# Non-ribosomal Peptides as Structural Determinants of Fungal Hydrophobicity

**DOI:** 10.64898/2026.07.13.738209

**Authors:** Trine Aalborg, Klaus R. Westphal, Balázs Delényi, Thea L. Lunden, Marie O. Jørgensen, Dmitry Tolmachev, Trine Sørensen, Maria Sammalkorpi, Jens L. Sørensen, Peter Kristensen, Herbert Kogler, Markus B. Linder, Reinhard Wimmer, Teis E. Sondergaard

## Abstract

Fungal surfaces must remain hydrophobic to enable growth, dispersal, and survival under fluctuating environmental conditions, yet the molecular basis of this property remains incompletely understood. Here, we identify fungisporins, fusahexins, and related cyclic non-ribosomal peptides (NRPs) as members of a conserved functional class of fungal metabolites, termed WAter Repellent Peptides (WARPs), that are required for fungal surface hydrophobicity. Across filamentous fungi, WARPs vary substantially in sequence and length but share conserved structural features, including cyclization, hydrophobic amino acid composition, and alternating D- and L-configurations, consistent with a flexible amphiphilic scaffold. Loss of WARP-producing non-ribosomal peptide synthetases results in rapid collapse of aerial hyphae upon water exposure, demonstrating that these peptides are required for maintenance of hydrophobic aerial structures. Using phage-display-derived antibodies, we localize WARPs to the hyphal surface, supporting their role as surface-associated structural components. Together, these findings identify a conserved NRPS-encoded peptide system that contributes to fungal hydrophobicity and establish WARPs as a broadly distributed class of surface-associated metabolites with structural function in filamentous fungi.

## 1 Introduction

A growing collection of orthologous, though highly diverged[1], non-ribosomal peptide synthetases (NRPSs) have been coupled to fungal surface hydrophobicity in five filamentous ascomycetes[1–7], and their manifold cyclic non-ribosomal peptide (NRP) products candidate a new addition to the group of universally occurring specialized metabolites (SMs). Here, we categorize these metabolites under the collective term WARPs (WAter Repellent Peptides). The described *warp* gene orthologues are *nrps4* in *Fusarium graminearum*[3,8], *hcpA* in *Penicillium chrysogenum*[4], *nrps2* in *Alternaria brassicicola*[5], *nrps4* in *Cochliobolus heterostrophus*[2], and *pes1* in *Aspergillus fumigatus*[6]. Deletion of the *warp* genes have been associated with distinct phenotypes, where mycelial colonies display an apparent aberration of surface hydrophobicity, losing their water repellant abilities[1–5]. Deletion mutants also present with altered mycelial surface structure[4], abnormal conidial cell wall morphology, reduced virulence, and compromised conidial germination rates[5]. Despite their prevalence, the NRPs have only been characterized for few of the *warp* genes, namely fungisporins from *Penicillium chrysogenum*[4] and *Aspergillus nidulans*[9] and fusahexin from *Fusarium graminearum[7]*. The ecological role and mode of function of the WARPs have hitherto remained elusive.

### 1.1 WARPs in *Ascomycetes*

Based on the known *warp* biosynthetic gene clusters (BGCs)[7], the canonical *warp* cluster comprises a *nrps* gene alongside an upstream ATP-Binding Cassette (ABC) transporter gene. The encoded NRPSs can have varying module counts, but as a rule every other module includes an epimerization domain (**Fig. S1, Tab. S1**). Upon reviewing fungal genomes within the NCBI database, the genetic ensemble responsible for the synthesis of WARPs are discernible in 5 out of 7 subfamilies encompassed by the predominant subgroup *Pezizomycotina* within *Ascomycetes*. Notably, numerous extensively researched fungi are featured within this panel, including representatives from *Sordariomycetes* such as *Chaetomium globosum* and all *Fusarium* species; *Eurotiomycetes* such as Aspergilli and all Penicillia; *Dothideomycetes* such as *Alternaria brassisicola* and *Cochliobolus heterostrophus*; as well as representatives from *Leotiomycetes* and *Lecanoromycetes*. It merits attention that all sequenced *Trichoderma* and *Arthrinium* species[10], lack a BGC conforming to the pattern of the *warp* BGCs. BGCs adhering to this pattern have not been identified in other fungi outside of the *Ascomycete* phylum, including basidiomycetes. Drawing from gene clusters identified in *Penicillium* and *Fusarium*, coupled with insights from this investigation, each NRPS may encompass between 3 and 6 modules, with the highest module count observed in *Apiospora*, totaling 6 modules, although the number of integrated amino acids may vary and can include iterative module application[7]. A discernible pattern also emerges in the peptide’s composition based on the alternating presence of epimerization domains in the NRPS modules and the identified WARPs, indicating the presence of alternating D*-* and L*-* enantiomers of largely hydrophobic amino acids.

## 2 Results

### 2.1 WARPs in *Aspergillus*

*Aspergillus* is one of the most extensively studied fungal genera, with many characterized secondary metabolites. It has long been known that they produce fungisporins[11,12], but the genetic biosynthetic origin of the compounds has not been definitively identified. It has previously been indirectly demonstrated that *nidA* is responsible for the production of fungisporins in *Aspergillus nidulans[9]*. Our sequence alignments with *Aspergillus fumigatus* indicate that fungisporins are synthesized by the gene *pes1*, which conforms to the modular organization of the WARP NRPSs and BGC structure (**Fig.1A**). To confirm this, we used a CRISPR/Cas9 approach to knock out the *pes1* gene in *As. fumigatus* (**Fig. S2**) and subsequently assessed the resulting phenotype (**Fig. 1B**) and fungisporin production (**Fig. S3**). It is challenging to perform drop tests and assess hydrophobicity in fungi with spore-covered surfaces, as spores are inherently water-repellent. To investigate the Δ*pes1* phenotype in *Aspergillus*, we cultured *As. fumigatus* mycelium mass in liquid culture under non-sporulating formative conditions followed by compression of dry mass for droplet testing. The knockout mutant displayed a distinctly phenotype wettable, directly comparable to those observed in *warp* deletion mutants of other fungal species (**Fig.1B**). Having identified the well-known phenotype, we proceeded to investigate the fungisporins produced by the Pes1 NRPS. The results showed that the dominant fungisporin was fungisporin C (cyclo[Tyr(D)-Tyr-Val(D)-Val]) (**Fig. S4**), conforming to the alternating D-/L-enantiomeric pattern observed for other WARPs.

**Fig. 1.**
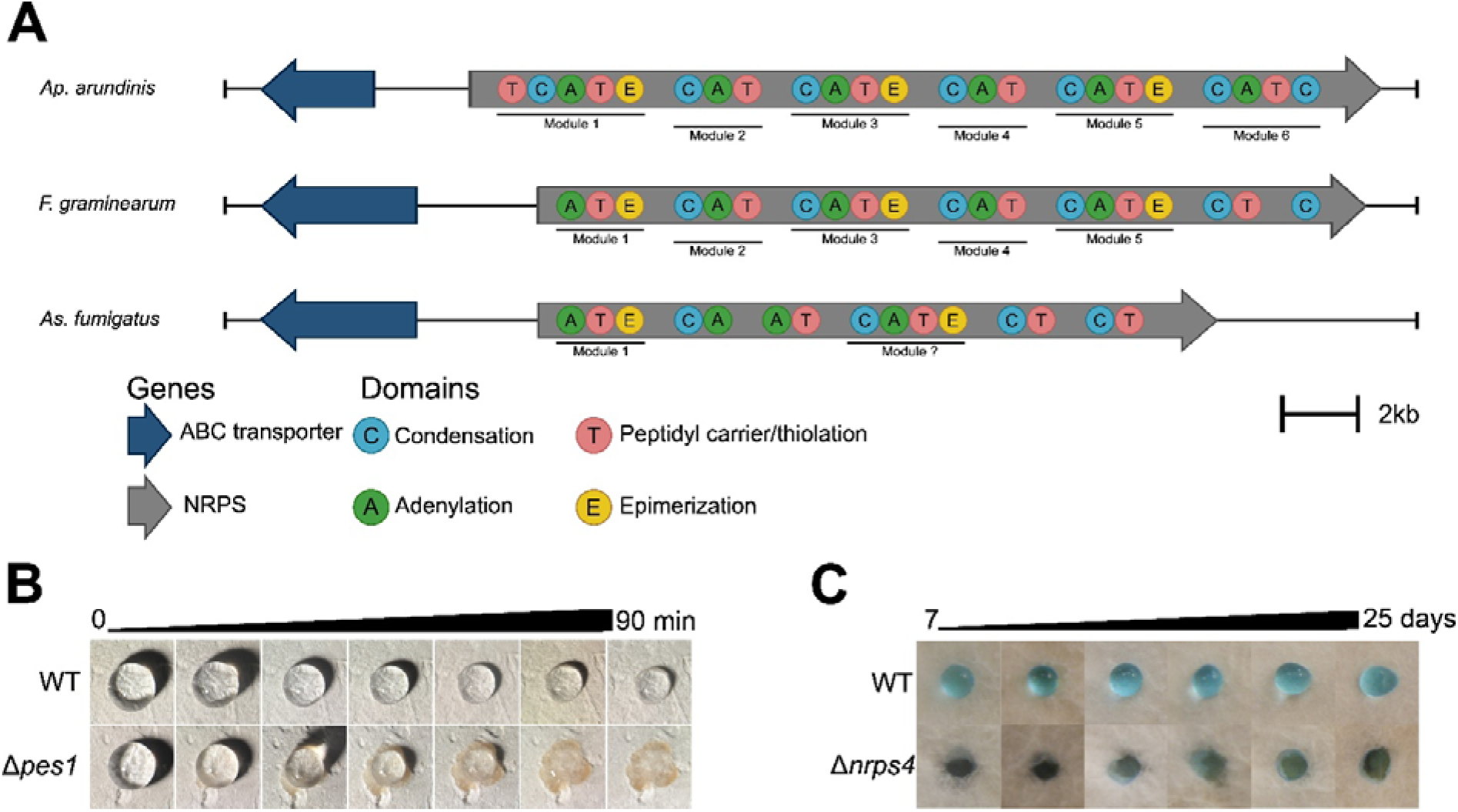
Products of the WARP NRPS biosynthetic gene cluster control fungal surface hydrophobicity. **A)** The *warp* BGC in *Ap. arundinis, F. graminearum*, and *As. fumigatus*. **B)** Water droplets are repelled on dried, pressed WT mycelium of *As. fumigatus* while the Δ*pes1* mycelium absorbs the droplet. Minutes indicate a time series of a single droplet absorption. **C)** Water droplets are repelled on vegetative WT *Ap. arundinis* AAU773 mycelium across a 25-day temporal period (individual droplets placed at each time point), while the Δ*nrps4* mycelium is completely penetrable.

### 2.2 WARPs in *Apiospora*

It is intriguing that two of the most extensively studied fungal genera, *Aspergillus* and *Penicillium*, produce WARPs containing four amino acids[4,9,11–15], while fusahexin from *Fusarium* is a cyclic hexapeptide, including a heterocyclic oxazine ring[7]. To explore the potential diversity of ascomycotal WARPs, we examined the *warp* gene in a species from the less-documented genus *Apiospora*[16]. Following isolation of a novel strain of *Ap. arundinis* (*Ap. arundinis* AAU773), whole-genome sequencing[17] revealed the *nrps4* gene as a propitious candidate locus for WARP synthesis, encoding a hexamodular NRPS fitting the consensus WARP NRPS module architecture (**Fig.1A**).

Employing a CRISPR/Cas9 approach, we targeted and disrupted the *warp* gene of *Ap. arundinis* AAU773 and verified mutants with long read sequencing (**Fig. S5**). Phenotypic analysis of the Δ*nrps4* mutant confirmed phenotypic characteristics comparable to those in other ascomycetes, where the vegetative mycelium of the wild-type (WT) remained completely water repellent over time and the Δ*nrps4* mutant displayed a total loss of surface hydrophobicity (**Fig.1C**).

### 2.3 Identification of the WARP apiosporin peptides

Comparative chemical analysis of the metabolite profiles of *Ap. arundinis* WT and Δ*nrps4* revealed that inactivation of the *nrps4* gene abolished biosynthesis of an array of compounds with related MS fragmentation pattern. A total of 14 different monoisotopic masses, some eluting as two distinct peaks on a hexyl-phenyl column, were absent from Δ*nrps4* extracts compared to WT (**Tab. S2, Fig. S6-S7**). The two most abundant masses were 814.5 [M+H]^+^ and 856.5 [M+H]^+^ (**Fig. 2A**). These were named apiosporin A and B, respectively. Four compounds with a mass-to-charge ratio of 814.5 [M+H]^+^ were obtained, along with two compounds with a mass-to-charge ratio of 856.5 [M+H]^+^.The compounds with the mass 814.5 [M+H]^+^ will from this point on be denoted as apiosporins A1 to A4 and the compounds with a mass of 856.5 [M+H]^+^ as apiosporins B1 and B2 (**Fig. 2B**). Structural elucidation by NMR demonstrated that apiosporins A1-A4 and apiosporins B1-B2 share identical molecular connectivities but differ in stereochemistry. Consistent with this, each stereoisomer displayed a distinct elution profile. Apiosporins A and B are cyclo-heptapeptides in which the lysine residue participates in formation of a heterocyclic 4-imidazolidinone ring. (**Fig. 2B**). The apiosporins A (cyclo[Leu-Leu(D)-Leu-Ala(D)-Leu-Tyr(D)-Lys]) and B (cyclo[Leu-Leu(D)-Leu-Leu(D)-Leu-Tyr(D)-Lys]) have varying composition of hydrophobic amino acids, indicating promiscuity of at least some A domains in addition to iterative module application, likely the first module 1. It is possible that in the TCATE module, the upstream T domain carries an L-Leu residue, while the A–T–E unit activates a second L-Leu, which is epimerized to D-Leu; the C domain then mediates condensation between the L- and D-Leu residues.

**Fig. 2.**
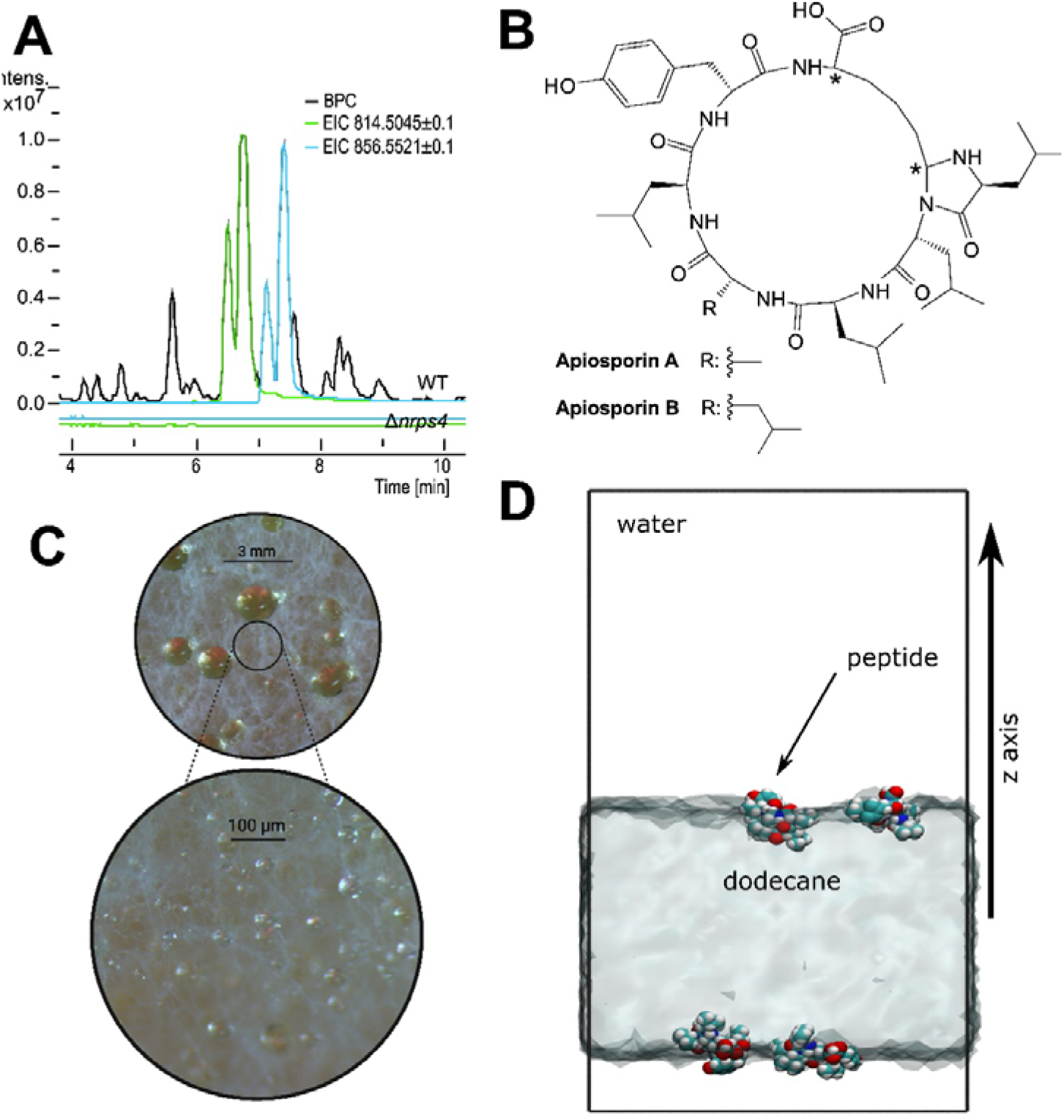
Apiosporins are hydrophobic cyclopeptides. **A)** Superimposed BPC (WT) and EICs of monoisotopic masses of apiosporins A (814.5 [M+H]^+^) and B (856.5 [M+H]^+^) in WT and Δ*nrps4* yeast-peptone-glucose (YPG) extracts before purification of A1-A4 and B1-B2 stereoisomers. **B)** Proposed molecular structures of apiosporins A and B. * Indicates stereochemical variation among apiosporin stereoisomers. **C)** Light microscope images of apiosporin-rich exudate droplets on the mycelial surface of *Ap. arundinis* AAU773. Δ*nrps4* did not produce exudate droplets on the mycelial surface. **D)** A visualization of molecular dynamics simulation of cyclo(Leu-Leu(D)-Leu-Ala(D)-Leu-Tyr(D)-Lys) peptides in a water/oil(dodecane) interface, with the explicit water molecules omitted in visualization and the dodecane slab positioning indicated by a continuum visualization for clarity.

(**Fig.1A**). The available structural data do not support iterative use of module 6. Instead, the terminal lysine-containing unit appears to participate in formation of the N,N-acetal-containing heterocycle, suggesting that module 6 is more likely involved in terminal incorporation and cyclization than in repeated elongation. Additionally, four of the monoisotopic masses absent from Δ*nrps4* extracts compared to WT corresponded to putatively cyclic and linear hexapeptide variants (**Tab. S2, Fig. S7**), further increasing the possible variability of the compounds produced by this single NRPS.

### 2.4 Extracellular localization of WARPs

We observed exudate droplets on the *Ap. arundinis* mycelial surface (**Fig. 2C**) and subsequently harvested and screened them for WARPs. Fungal guttation or exudate droplets are commonly encountered in various fungi, primarily triggered by fluctuations in temperature, salinity, or nutrient availability, leading to alterations in osmotic or turgor pressure[18]. Several SMs, lipids, enzymes, and proteins have been identified within fungal exudate droplets[19], and our analysis revealed that the predominant compounds in *Ap. arundinis* exudates include apiosporins A and B (**Fig. S8**), indicating that these WARPs are located extracellularly. Extracellular localization of the WARPs is not surprising, as all annotated *warp* BGCs contain an ABC transporter gene (**Fig. S1**) and fungisporin has been identified in extracellular vesicles in *Penicillium digitatum*[20]. Based on molecular dynamics simulated interface systems, WARPs can spontaneously orient at aqueous–nonpolar interfaces (**Fig. 2D**). In fungi, such peptides may localize at or beyond the cell wall, where complex amphiphilic environments exist. The hydrophobic microdomains of the cell wall and the amphiphilic nature of extracellular exudate droplets provide suitable interface for these molecules, supporting their extracellular accumulation and surface-associated functions. To localize the apiosporins, we used the Predator single domain phage display library[21] to select antibodies specifically binding apiosporin (**Fig. S9**). Staining using these antibodies showed that the apiosporins are distributed evenly across the hyphal surface (**Fig. 3, Fig. S10**).

**Fig. 3.**
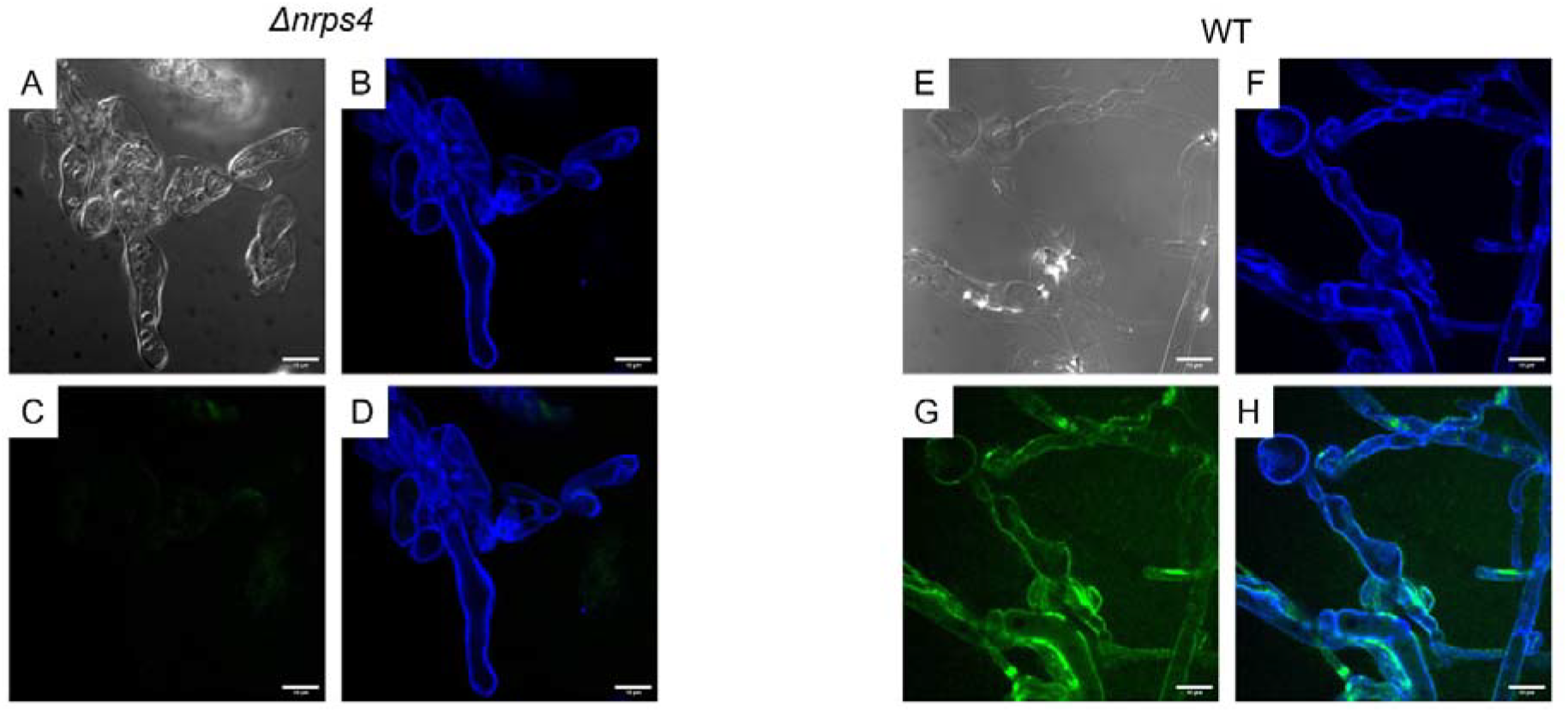
Apiosporins are embedded in the fungal cell wall based on selective staining. Selective staining for apiosporin A in the Δ*nrps4* and *Ap. arundinis* AAU773 (WT) with an Alexa 488 marked selected single domain secondary antibody [AP2] (green) and general staining of the fungal cell wall with Calcofluor white stain (blue). Δ***nrps4*: A)** Light microscopy, **B)** Calcofluor white, **C)** AP2, **D)** Combined calcofluor white and AP2. **WT: E)** Light microscopy, **F)** Calcofluor white, **G)** AP2, **H)** Combined calcofluor white and AP2. Size bar = 10 µm.

### 2.5 Molecular dynamics simulations of WARPs demonstrate conserved amphiphilic properties

WARPs show high structural variability (tetra-to heptapeptides), suggesting diverse properties. Molecular dynamics simulations of the smallest (tetra-) and largest (hepta-) peptides (**Fig. S10**) in bulk water and at a model oil/water interface reveal disparate flexibility and aggregation in water, yet all localized at the interface, indicating amphiphilicity. Sequence-dependent hydrophobicity affected partitioning, with more hydrophobic peptides positioned deeper in the oil phase. This interfacial localization is consistent with association with hydrophobic–hydrophilic interfaces in fungal surface environments. Both pure L- and alternating L-/D-heptapeptides showed similar amphiphilic behavior, suggesting that interfacial activity stems from structural rather than stereochemical factors; stereochemical heterogeneity likely provides proteolytic resistance[22– 24]. Cyclization similarly enhances stability[25]. Heptapeptides exhibited greater flexibility than tetrapeptides due to longer backbones and heterocyclization with the flexible Lys side chain, allowing more favorable interactions and stronger aggregation responses, while single-residue changes had proportionally larger effects in tetrapeptides.

## 3 Discussion

### 3.1 One NRPS can synthesize a collection of structurally similar WARPs

Predicting NRP structures from NRPS modular organization is difficult. While most NRPSs follow a linear mechanism with an initiation (A–T) and several elongation (C–A–T) modules, some act in iterative or non-linear ways, breaking the colinearity rule, where module number does not match peptide length[26]. The four-module HcpA NRPS of *P. chrysogenum* produces the hydrophobic, cyclic tetrapeptide fungisporin A (cyclo[Phe(D)-Phe-Val(D)-Val])[4], originally isolated in 1952 from the spores of several species of both *Penicillium* and *Aspergillus*[27,28]. In addition to fungisporin, HcpA also synthesizes at least nine other variants of the compound with differing amino acid compositions; its four adenylation domains are able to promiscuously recognize Phe/Tyr, Phe/Trp, Val/Ile, and Val/Ile, respectively[4]. The five-modular NRPS4 of *F. graminearum* produces fusahexin (cyclo[Ala(D)-Leu-*allo*-Thr(D)-Pro-Leu(D)-Leu]), a cyclic hexapeptide with a rare oxazine heterocyclic ring. Notably, fusahexin includes one additional amino acid compared to the number of A domains harbored by NRPS4, thought to originate from the iterative usage of one module facilitated by a C-terminal C-T-C tridomain[7] (**Fig. 1A**). We reinvestigated the produced fusahexins from *F. graminearum* and found several hydrolyzed variants[7]. In our study of *Ap. arundinis*, at least eight cyclic and two putatively linear apiosporin heptapeptides as well as two cyclic and two putatively linear apiosporin hexapeptides have been identified (**Tab. S3**). The diverse array of WARP variants can hence be produced by 1) hydrolysis, converting them into linear forms, 2) promiscuous adenylation domains capable of integrating a selection of physicochemically similar, but distinct amino acids, and 3) selective iterative application of adenylation domains, allowing deviation from the collinearity rule and production of WARPs of various lengths or, alternatively, proteolysis. There are several potential reasons for the multifarious WARP product profile found in each species, e.g. intentional utilization as stabilizers or adhesives between individual hyphae, where integration of a motley compound pool might lead to either varying interactions in the cell wall or a mutable cell wall profile in response to environmental cues. It could also be a result of evolutionary adaptation of an essential SM group to the broad niche-range of *Ascomycetes*, to adopt a substrate-permissive synthesis pathway. Characterizing the fungal WARPs is cumbersome, as it involves identifying not a singular substance per species but rather a series of related compounds with chemical modifications, diverse amino acid compositions, and complex stereochemistry. Though structurally distinct, these related compounds may fulfill the same biological function in their respective host organisms, despite substantial variation in amino acid composition, stereochemistry, and peptide length. Two other compound series have also previously been hypothesized to be products of *Fusarium* NRPS4 homologues[7,8], namely the lipo-peptides acuminatum A-C (cyclo[3*S*,4*R*-HMTA-*allo*-Thr(D)-Ala-Ala(D)-Gln-Tyr(D)-Leu/Ile/Val])[29] and a collection of nine unnamed hexacyclopeptides[30] (cyclo[Hyp/Dhp-Xle-Ile/Leu-Ala/Val-Thr-Xle]), though their ecological role has not been determined. Furthermore, malpibaldins A-C are cyclo-pentapeptide biosurfactants (cyclo[Val(D)-Leu-Leu(D)-Phe/Trp/Tyr(D)-Leu]) that have been isolated from the zygomycete *Mortierella alpina*[31–33] with similar molecular structures to WARPs. Bioinformatic analysis revealed two putative NRPSs with modular organization compatible with the biosynthesis of the NRPs (MpbA and MpcA)[33], also matching the suggested canonical WARP NRPS modular structure with epimerization domains in every other module. Based on the observed diversity of WARPs produced by individual synthetases and their shared association with fungal surface hydrophobicity, we propose an ONMAC (One NRPS Many Compounds) principle in which a single NRPS generates a family of structurally related metabolites that collectively fulfill the same biological function. **Table S4** holds a list of known and putative WARP compounds and their producing species. We suggest classifying the WARPs by the number of amino acid residues.

### 3.2 Surface hydrophobicity and the WARP hypothesis

Surface hydrophobicity in filamentous fungi has widely been attributed to hydrophobins, a group of small amphipathic proteins found in the outermost layer of the cell wall of hyphae and spores[34,35]. The spatial structure of hydrophobins logically supports this role, as one face of the protein exposes aliphatic sidechains forming a large hydrophobic patch[36]. For class II hydrophobins, structural analysis and modelling show how the proteins can self-assemble into a two-dimensional hexagonal pattern with each hydrophobic patch exposed to one side[37]. One can easily picture how an interlocked film of these multimeric protein complexes can render the fungal surface hydrophobic. Class I hydrophobins are suggested to have another arrangement in which they form amphiphilic rodlets[38]. The role of hydrophobins in mycelial surface hydrophobicity is supported by deletion strains that show easily wettable phenotypes[39]. However, phenotypes are often only marginally more wettable, indicating more complex mechanisms. For example, in *F. graminearum* the deletion of all hydrophobin genes did not result in wettable phenotypes[40]. Similar results have been obtained with *Botrytis cinerea[41]* and *Metarhizium robertsii[42]. In vitro* studies of hydrophobin physicochemical properties are challenging since hydrophobins can be very difficult to isolate and perform mechanistic studies on. However, studies have reported that hydrophobins can increase the hydrophobicity of various surfaces *in vitro*[43,44]. Given the close correspondence in ring architecture, it seems reasonable that our naturally occurring cyclic peptide could engage in the same antiparallel stacking and backbone-directed hydrogen-bonding interactions described by Ghadiri et al. (1993)[45], and thus adopt comparable tubular assemblies.

Our hypothesis for integration of WARPs in the fungal cell wall is shown in **Fig. 4**. Inactivation of both WARPs and hydrophobins yields similar phenotypes, suggesting a shared role in forming the fungal cell wall’s hydrophobic “molecular raincoat.” Hydrophobins occur in spores, likely with melanin forming a water-repellent layer, though their presence in hyphae is less clear. Water repellence thus involves more than hydrophobins alone. We propose that both WARPs and hydrophobins contribute to hydrophobicity, with strain-specific combinations reflecting their diversity and potential synergies. Class II hydrophobins occur only in Ascomycetes, Class I in Ascomycetes and Basidiomycetes, whereas WARPs are confined to Ascomycetes. The stronger hydrophobic defect observed in WARP mutants compared to reported hydrophobin mutants raises the possibility that WARPs contribute centrally to fungal surface hydrophobicity, while hydrophobins may provide complementary or partially overlapping functions[40]. Cell wall anchoring mechanisms remain unclear but are likely to involve non-covalent crosslinking[46,47]. Disrupting this proposed network causes major phenotypic effects, as seen for both hydrophobin and WARP deletions.

**Fig. 4.**
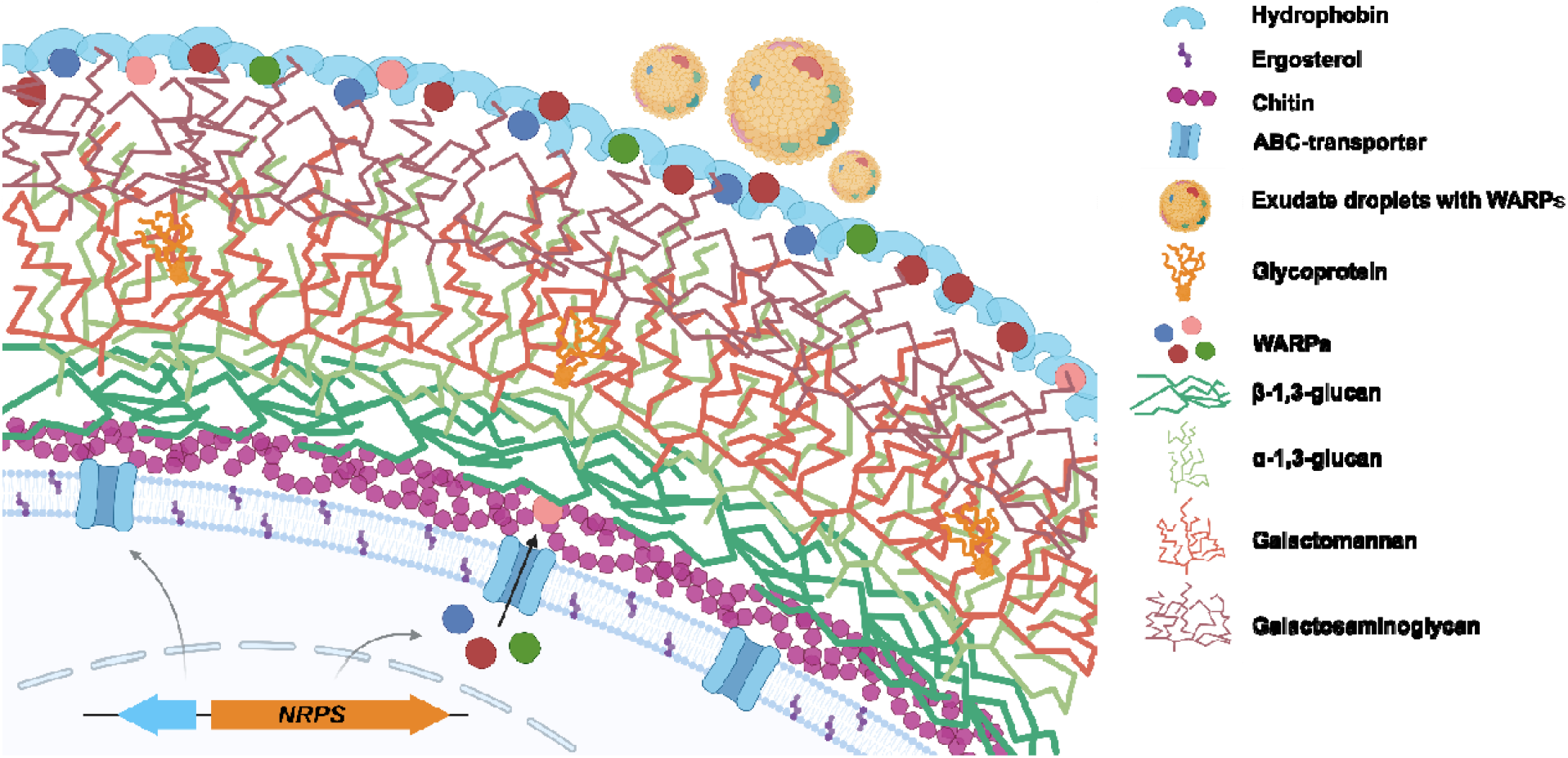
Proposed structure of the fungal cell wall including WARPs. Proposed schematic model of the fungal cell wall in *Ascomycetes*. A battery of structurally related compounds (WARPs) is produced intracellularly at least in part by an NRPS and is exported to the extracellular space by a cluster-associated ABC transporter. The WARPs are integrated into the cell wall, hypothetically by non-covalent interactions in the hydrophobin film on the outward mannoprotein layer and function to mediate the water-repellent surface properties of the cell wall in association with the hydrophobins – this mosaic of multifarious hydrophobins and WARPs constitutes a fungal raincoat. WARPs are incorporated into lipid droplets during the guttation of high-affinity fatty acids. Hyphal cell wall composition was based on *As. fumigatus*[48].

A bipartite mechanism for hydrophobicity may offer ecological advantages. NRPS-mediated WARP production enables rapid, energy-efficient synthesis and flexible regulation. In *Fusarium*, aerial mycelia lose water repellence after 10–14 days, but WARP overexpression maintains it[3], aligning with aerial mycelium collapse during perithecia formation described by Hein (1928)[49]. WARPs may reinforce vegetative hyphae, while stationary hyphae and spores rely on other compounds and hydrophobins[50]. For example, in the yeasts spore outer cell wall layer contains interconnected LL and DL-dityrosine to increase proteolysis and thermal resistance of the spores, but not the vegetative cell wall[51].

Our findings add WARPs to a growing list of fungal metabolites that contribute directly to cellular architecture rather than serving exclusively ecological, defensive, or virulence-related functions. While melanin has long been recognized as a structural cell-wall component, recent studies have identified additional specialized metabolites required for rodlet-layer formation and cell-wall semipermeability in pathogenic fungi[52,53]. WARPs extend this emerging paradigm to a conserved family of NRPS-derived cyclic peptides distributed across filamentous Ascomycetes. Despite substantial variation in amino acid composition, stereochemistry, and peptide length, WARPs are united by a common biological function associated with fungal surface hydrophobicity. Their extracellular localization, conserved amphiphilic properties, and the wettable phenotype observed upon loss of WARP biosynthesis support a role in maintaining the integrity of hydrophobic aerial structures. More broadly, these findings suggest that specialized metabolism is not restricted to ecological interactions, chemical defense, or virulence, but can also contribute directly to the physical construction and functional organization of the fungal body. We propose that WARPs represent a specialized metabolite system integrated into fungal surface architecture and provide a framework for understanding how structurally diverse metabolites can converge on a common structural function.

## Resource availability

Study data are included in the article.

## Acknowledgements

Computational resources by CSC IT Centre for Finland and RAMI -- RawMatters Finland, are acknowledged. We acknowledge CSC for awarding this project access to the LUMI supercomputer, owned by the EuroHPC Joint Undertaking, hosted by CSC (Finland) and the LUMI consortium through a CSC project call.

## Supplemental information

**Document S1: Figures S1-S21, Tables S1-S16, Supplementary references for Table S4**.

