## Supplementary material for "Non-ribosomal Peptides as Structural Determinants of Fungal Hydrophobicity": Sup

### Supplementary Figures

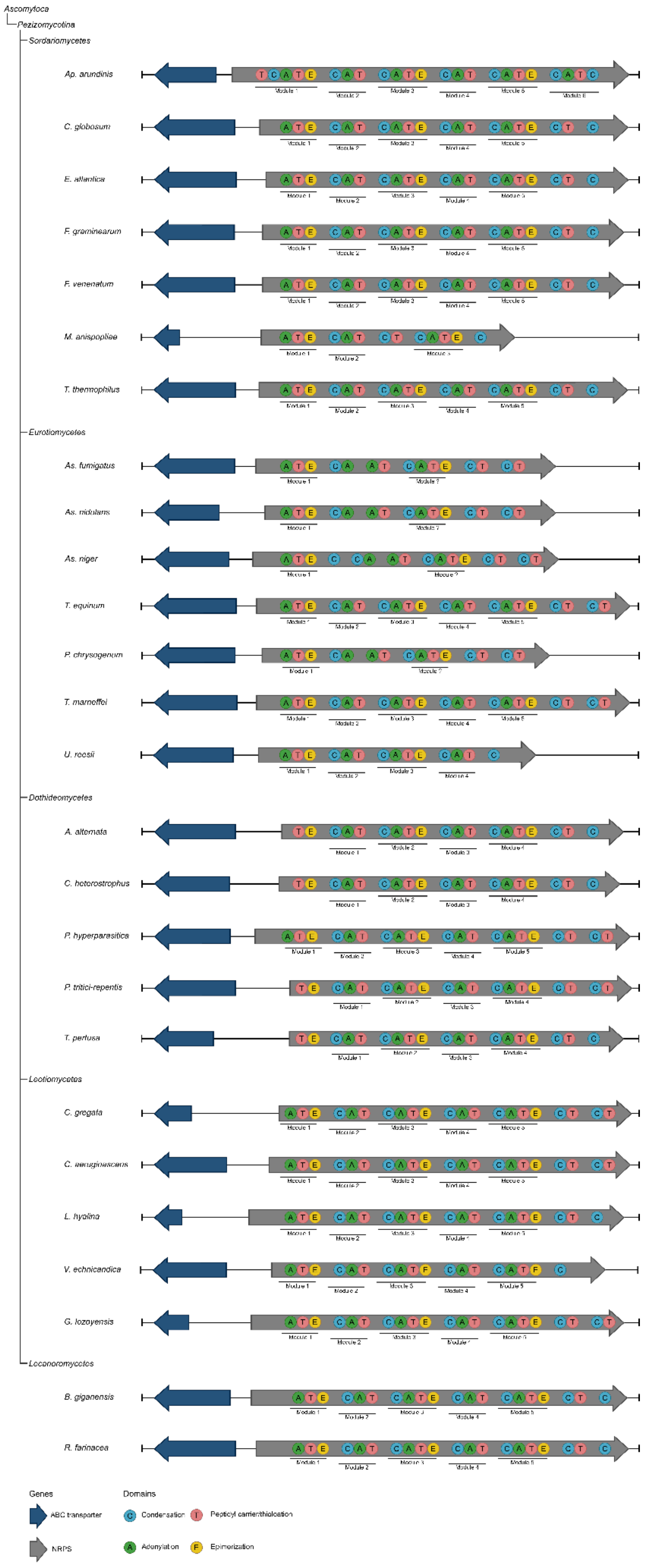
Ascomycota
Pezizomycotina
Sordariomycetes
Ap. arundinis
C. globosum
E. atlantica
F. graminearum
F. venenatum
M. anisopliae
T. thermophilus
Eurotiomycetes
As. fumigatus
As. nidulans
As. niger
T. equinum
P. chrysogenum
T. marneffei
U. reesii
Dothideomycetes
A. alternata
C. heterostrophus
P. hyperparasitica
P. tritici-repentis
T. pertusa
Leotiomycetes
C. gregata
C. aeruginascens
L. hyalina
V. echinocandica
G. lozoyensis
Lecanoromycetes
B. gigantensis
R. farinacea
Genes
ABC transporter
NRPS
Domains
Condensation
Peptidyl carrier/thiolation
Adenylation
Epimerization

Fig. S1. Phylogeny of WARP biosynthetic gene clusters across Ascomycota. WARP BGCs from representative species of the subgroup Pezizomycotina within the phylum Ascomycota are shown, with taxonomic classes indicated by the tree-structure on the left. The clusters exhibit highly conserved gene architectures, including a neighbouring ABC transporter. Predicted modules and domains of the NRPS are displayed to highlight the conserved organization, and gene sizes are shown to scale relative to one another. Species and corresponding gene identifiers are provided in Tab. S1.

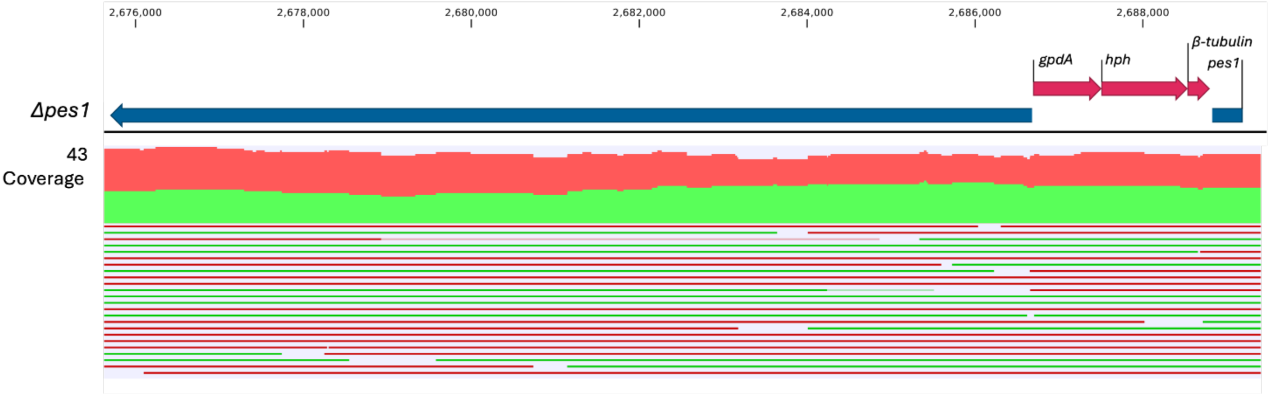

Fig. S2. Whole-genome sequencing As¡.fumigatus Δpes7 mutant validation. Full genome sequencing of the As¡.fumigatus.Δpes7 mutant. The mutant was generated by CRISPR/Cas9 cloning and verified by long read Nanopore sequencing. Reads mapped to HygR resistance cassette insert of modified As¡.fumigatus genome demonstrates correct insertion of the cassette.

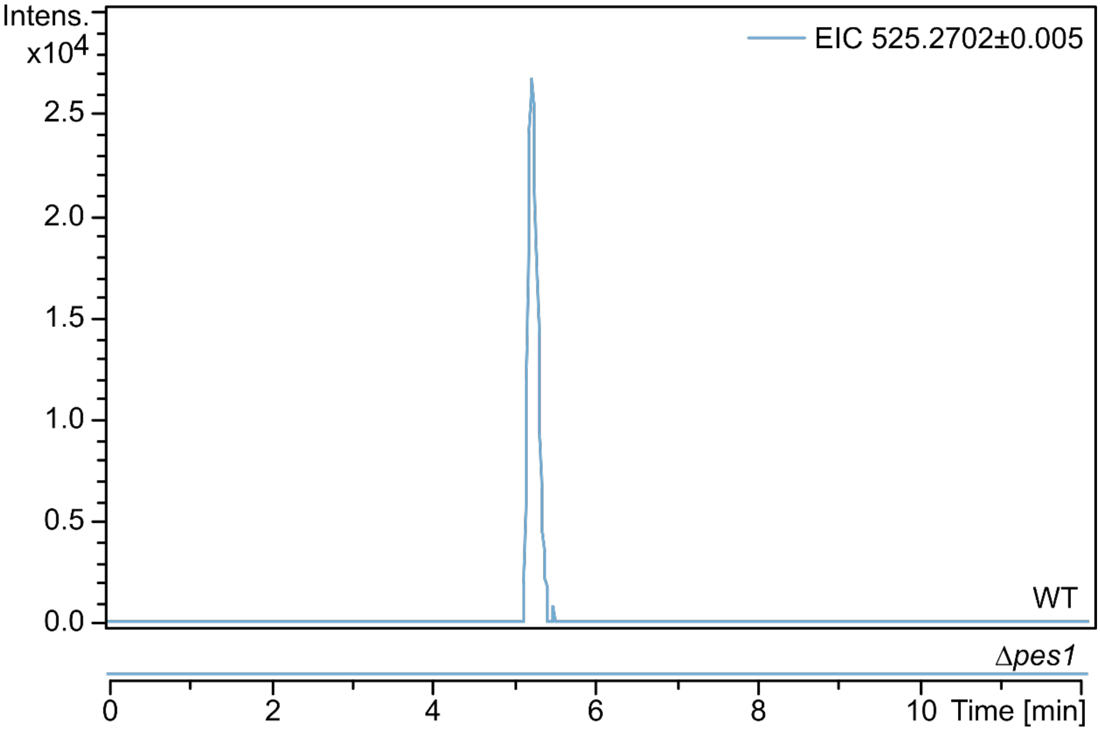

Fig. S3. LC-MS of fungisporin C in As¡.fumigatus WT and Δpes7. Extracted ion chromatograms (EICs) of harvested As¡.fumigatus WT and Δpes7 CYA microextracts and of Pes1-derived fungisporin C monoisotopic mass ($[M+H]^+$ = 525.3).

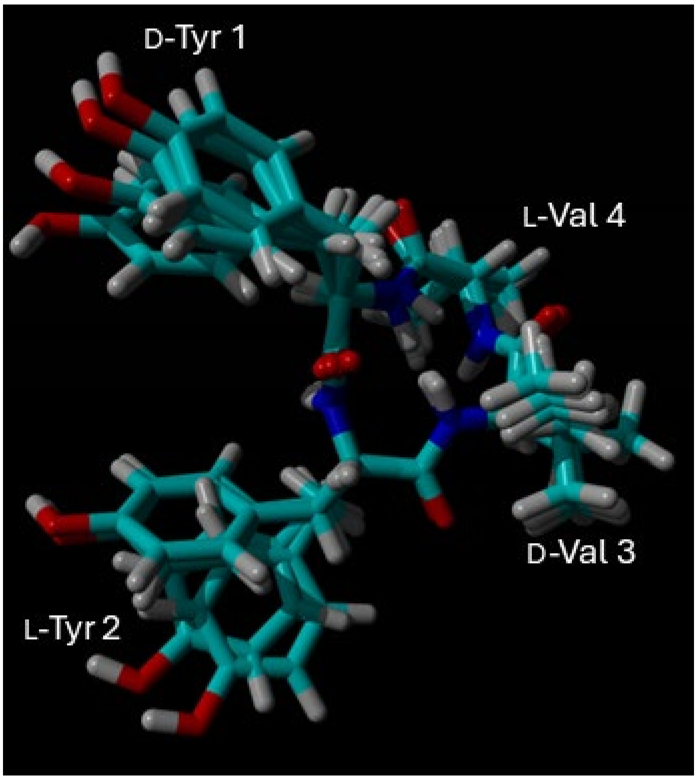

Fig. S4. Compound structure of fungisporin C. A possible structure of cyclo(Tyr(D)-Tyr-Val(D)-Val) energy-minimized with all $\phi$= -120° ±25°. Bundle of four energy-minimized structures of fungisporin with $\phi$= -120° ±25°, as derived from NMR data. Structures were superposed on backbone N, $C^{\alpha}$ and C' atoms.

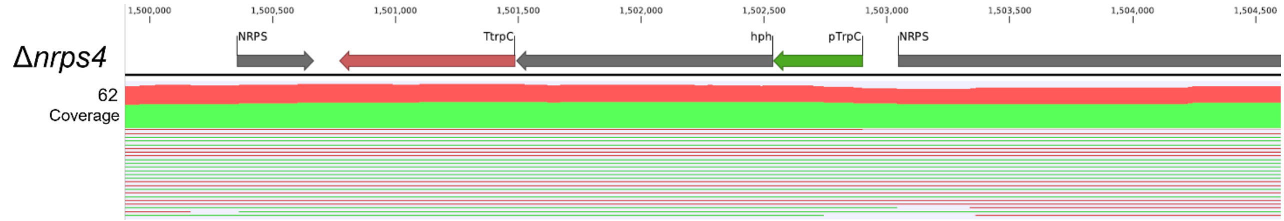

Fig. S5. Whole-genome sequencing Ap¡.arundinis Δnrps0 mutant validation. Full genome sequencing of the Ap¡.arundinis AAU773 Δnrps0.mutant. The mutant was generated by CRISPR/Cas9 cloning and verified by long read Nanopore sequencing. Reads mapped to HygR resistance cassette insert of modified Ap¡.arundinis AAU773 genome demonstrates correct insertion of the cassette.

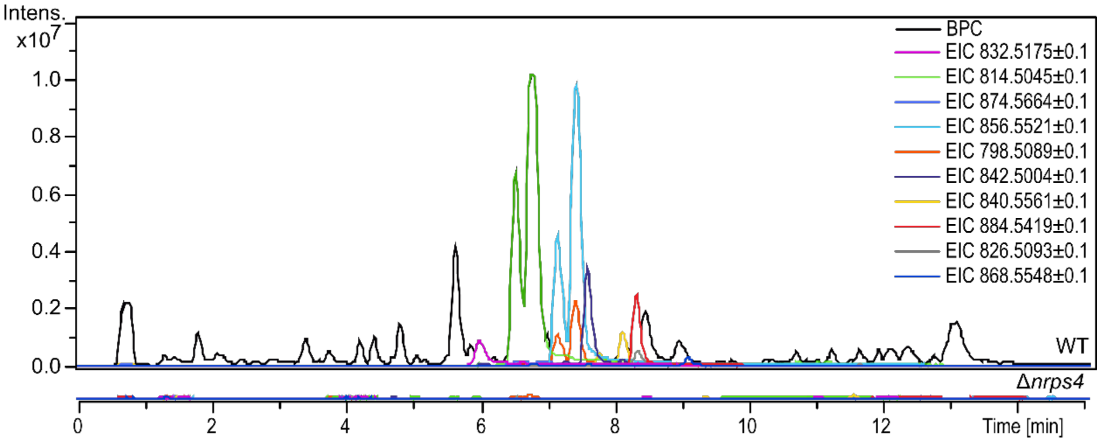

Fig. S6. LC-MS of Ap¡.arundinis heptapeptide apiosporins in WT and Δnrps0¡ Superimposed base peak chromatogram (BPC) of harvested Ap¡.arundinis AAU773 WT and Δnrps0 YPG microextracts and extracted ion chromatograms (EICs) of heptapeptide monoisotopic masses absent from Δnrps4 extracts.

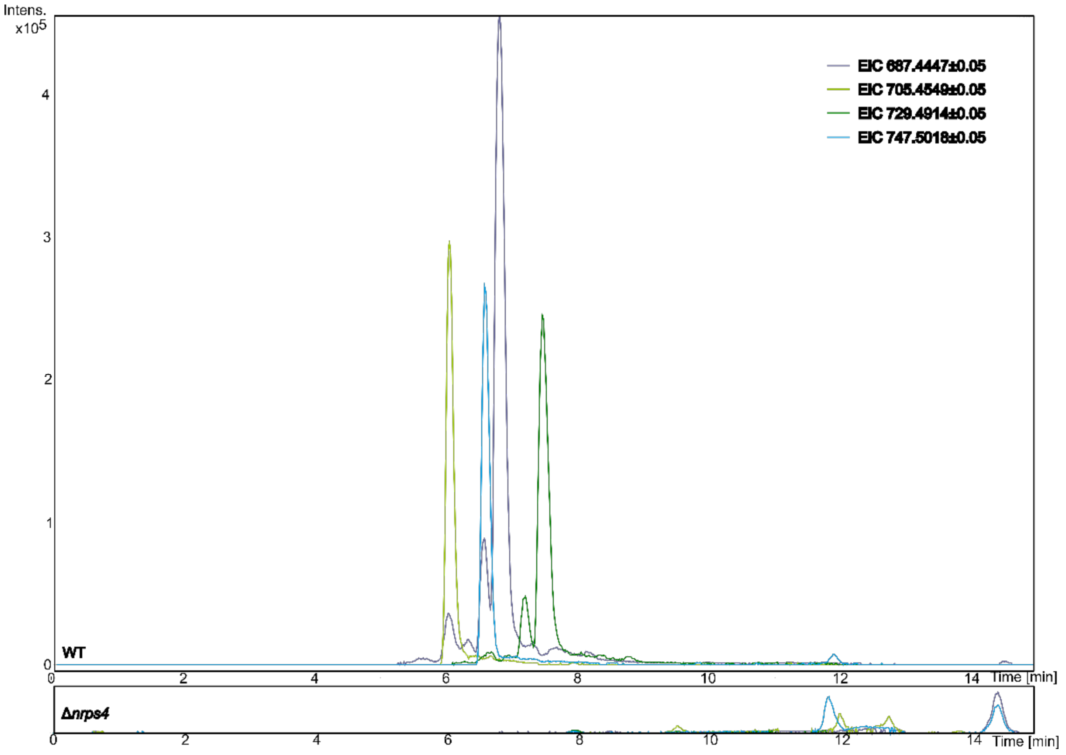

Fig. S7. LC-MS of Ap¡.arundinis hexapeptide apiosporins in WT and Δnrps0. Extracted ion chromatograms (EICs) of harvested Ap¡.arundinis AAU773 WT and Δnrps0 YPG microextracts corresponding to hexapeptide monoisotopic masses absent from Δnrps4 extracts.

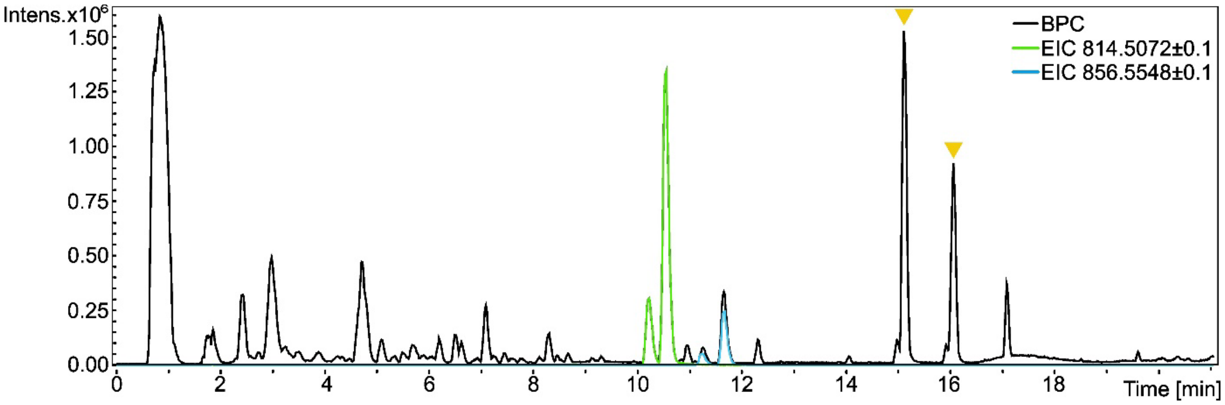

Fig. S8. LC-MS of apiosporins A and B in exudate droplets of Ap¡.arundinis WT. Superimposed base peak chromatogram (BPC) of harvested Ap¡.arundinis AAU773 WT lipid droplets and extracted ion chromatograms (EICs) of [Apiosporin A + H]$^+$ ± 0.1 and [Apiosporin B + H]$^+$ ± 0.1 monoisotopic masses. Yellow arrow heads indicate monoisotopic mass peaks of putative lipids.

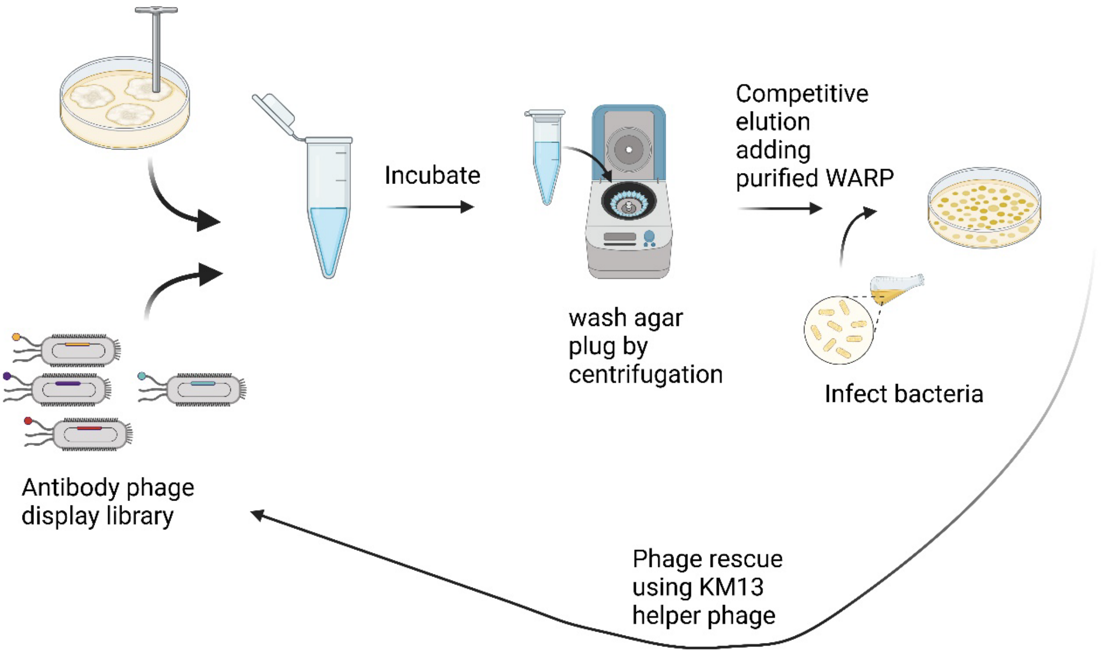

| |
|---|
| Sequence AP2:<br>EVQLLESGGGLVQPGGSLRLSCAASGFRDSDEDMGWVRQAPGKGLEWVSSIETPDGSTYYADSVKGRFTISRD<br>NSKNTLYLQMNSLRAEDTAVYYCASQPVYSYHFDYWGQGTLVTVSS |

Fig. S9. Overview of Phage display method and the sequence of AP2.

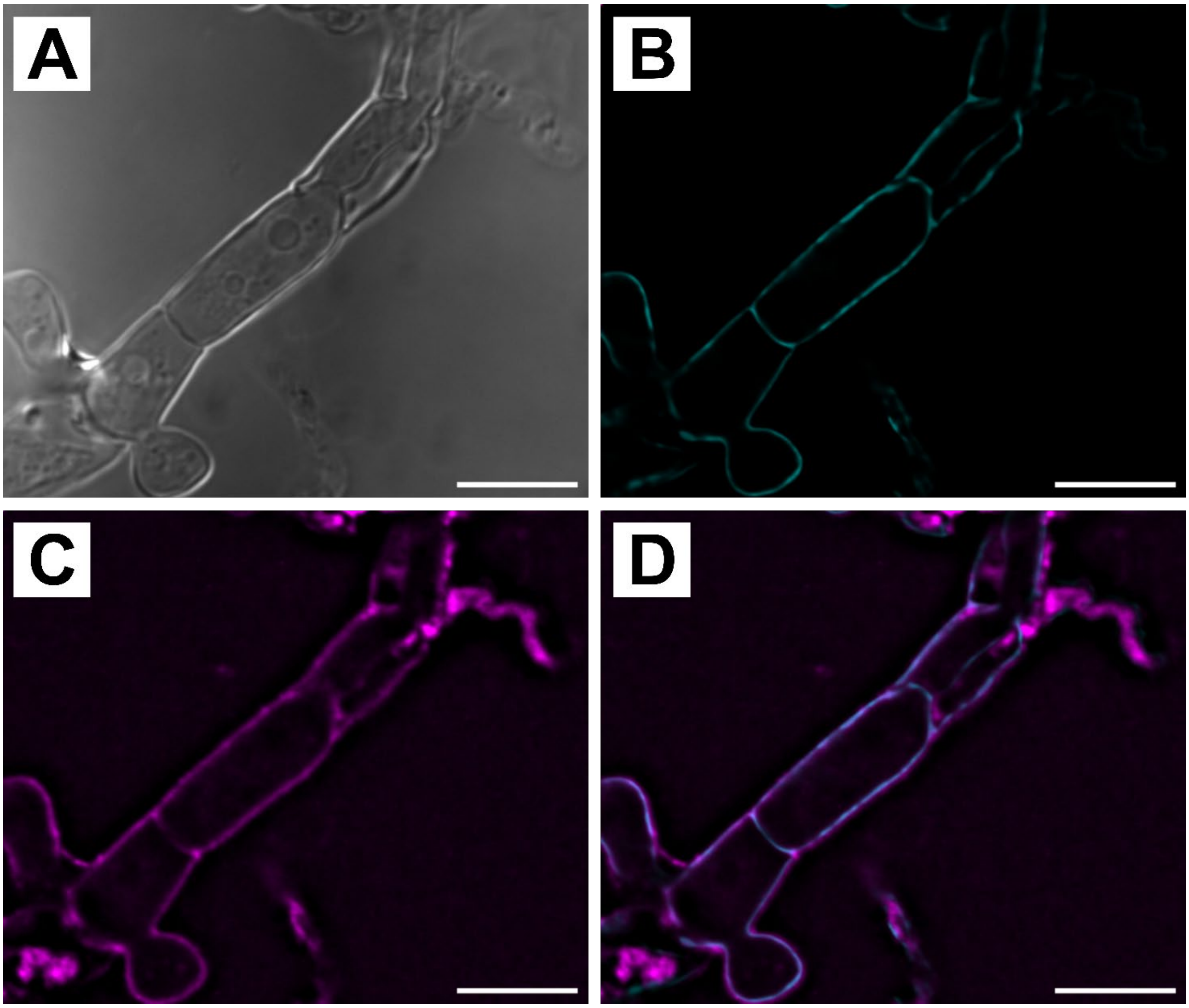

Fig. S10. Apiosporins localize to the hyphal surface based on selective antibody staining. Selective staining for apiosporin A in the Ap¡.arundinis AAU773 (WT) with an Alexa 488 marked selected single domain antibody [AP2] (magenta) and general staining of the fungal cell wall with Calcofluor white stain (cyan), Images were processed by deconvolution. A) Bright-field image B) Calcofluor white, C) AP2 staining D) Merged image. Size bar = 10 μm.

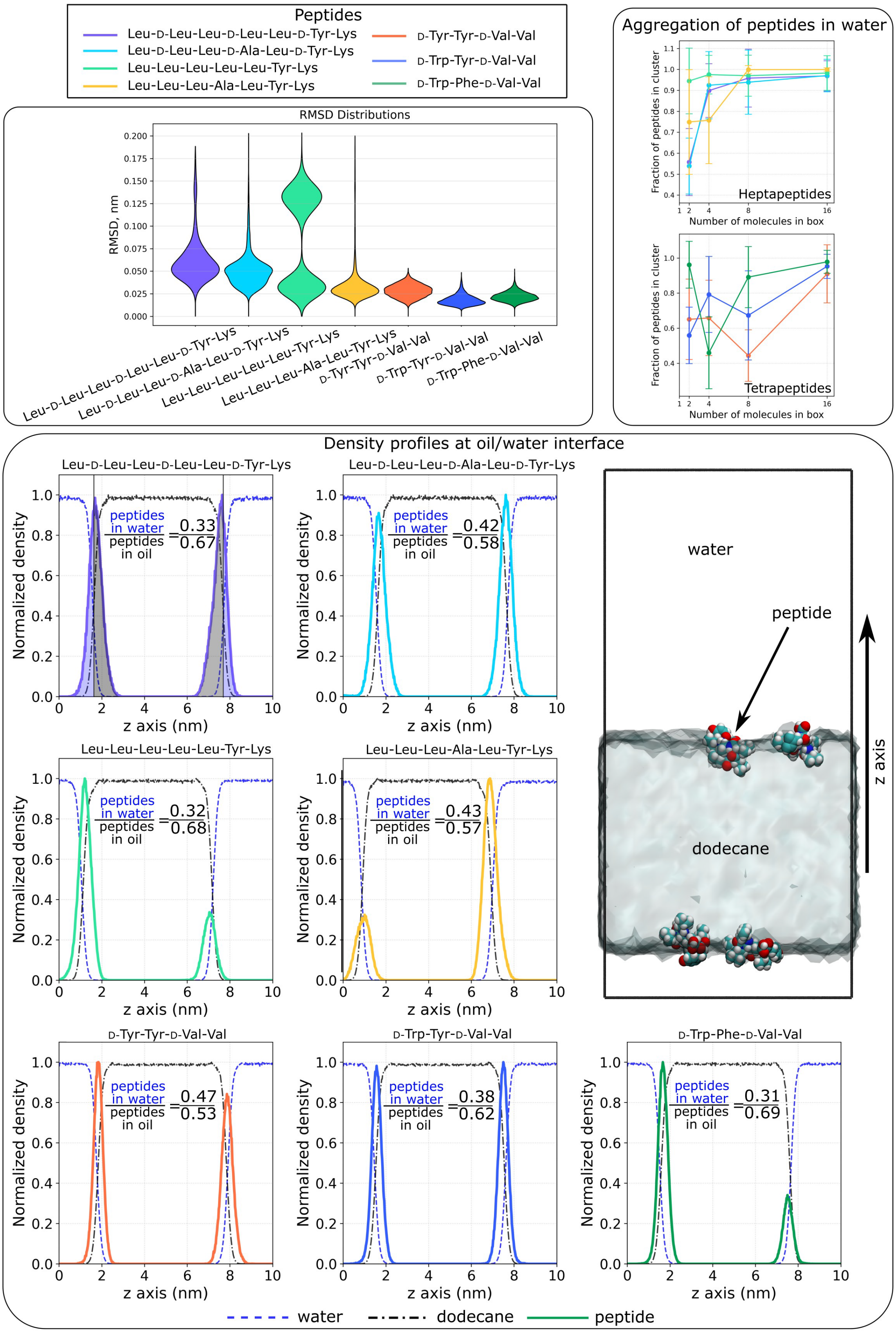

Fig. S11. Molecular Dynamics simulations results. Since the fungal cell wall consists of a large number of components, such as polysaccharides, glycoproteins, and hydrophobins, full scale atomistic simulation of this structure even at model level is challenging.  Therefore, the water/oil interface was used as a simplified model of the interphase between two media with different polarities (methods section in the SI). Top: Backbone RMSD distributions for individual peptides in water (left) and the fraction of clustered peptides at different peptide concentrations in water (right). The clustering is assessed based on the largest cluster. Bottom: Normalized number-density profiles calculated perpendicular to the oil/water interface (z-axial density profiles). Dashed lines correspond to water (blue) and dodecane (black) whereas solid lines show the peptide distributions. In all systems, the peptides reside at the interface. Inset fractions show peptide partitioning ratio between water and oil. Example of the partitioning ratio integration shown for the cyclo(Leu-Leu(D)-Leu-Leu(D)-Leu-Tyr(D)-Lys) system. A visualization of the simulated interface system for the system with 4 cyclo(Leu-Leu(D)-Leu-Ala(D)-Leu-Tyr(D)-Lys) peptides, with the explicit water molecules omitted in visualization and the dodecane slab positioning indicated by a continuum visualization for clarity.

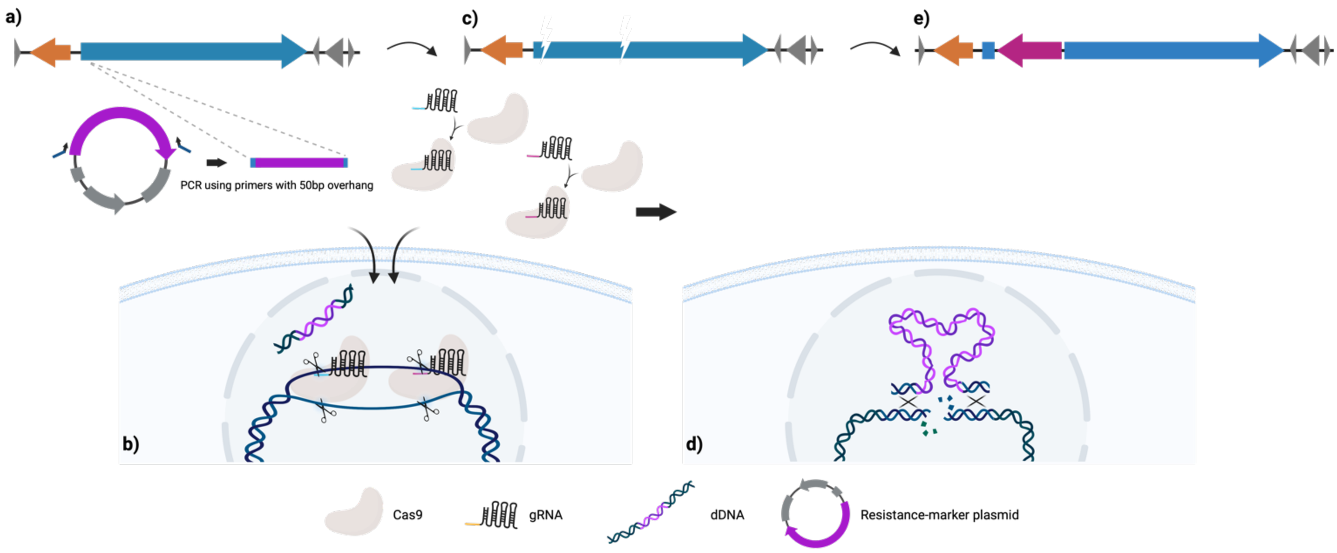

Fig. S12: Schematic of CRISPR/Cas9 genome editing approach in Aspergillus.fumigatus. a) A dDNA cassette containing the HygR resistance marker is amplified by PCR. The primers used have 50 bp overhangs matching the homology regions at the target exon 1 and exon 3 cut-site, creating microhomology arms for homology-directed repair. b) The dDNA cassette and in.vitro assembled RNA complexes of crRNA::tracrRNA and Cas9, targeting exon 1 and exon 3 respectively, are transformed into protoplast of the fungus by PEG-mediated transformation. c) The RNP complexes introduces a double-stranded break at the respective target site in the genome. d) The insertion fragment of the dDNA cassette is integrated into the genome at the target cut-sites by homology-directed repair facilitated by the 50 bp cut-site microhomology arms of the dDNA cassette resulting in a deletion of 7,802 bp. e) The pes7 gene is inactivated by deletion of 7,802 bp in addition to stable integration of the selection marker into the genome.

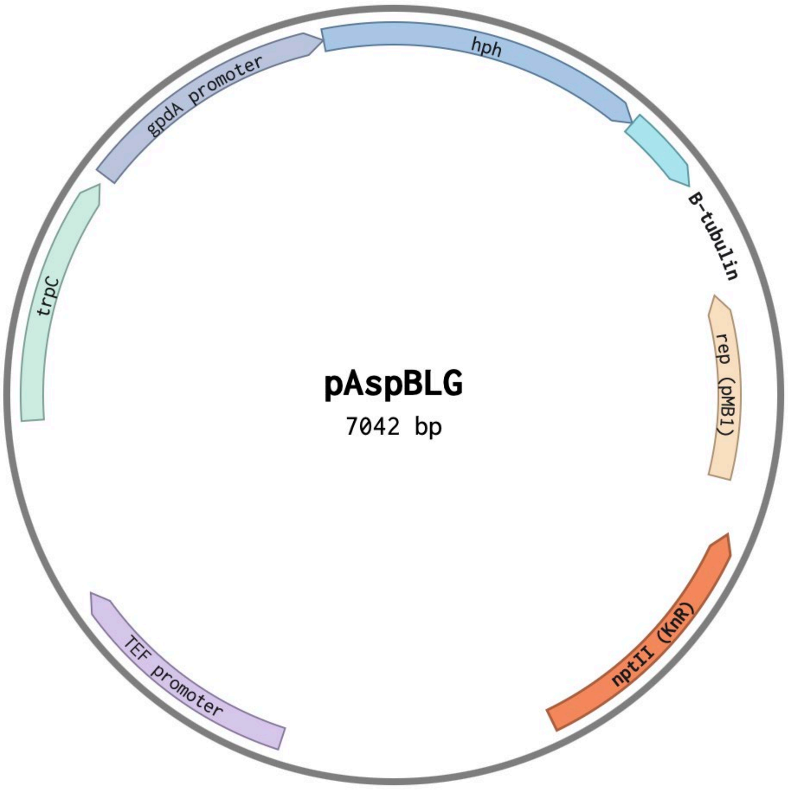

Fig. S13: Plasmid map of the pAspBLG vector.

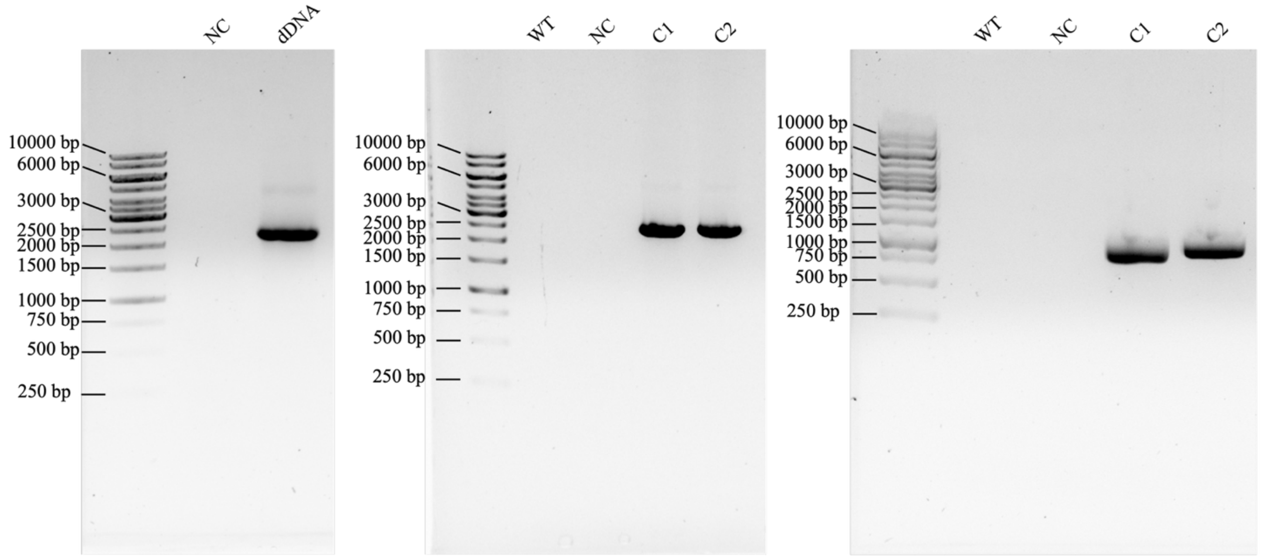

Fig. S14: Agarose gels (1%) of dDNA and colony screen for *As. fumigatus* Δ*pes1* mutant generation. Left: dDNA amplified with 50 bp overhang primers. Middle: positive colonies are screened using primers positioned inside and outside the insert (primer 3 and 4). A correct insertion produces a fragment of the expected size 2,305 bp. Left: positive colonies are screened using primers positioned inside and outside the insert (primer 5 and 6). A correct insertion produces a fragment of the expected size 997 bp. WT refers to wildtype, C1 and C2 refers to colony 1 and colony 2, respectively, and NC is a negative control without DNA template for PCR.

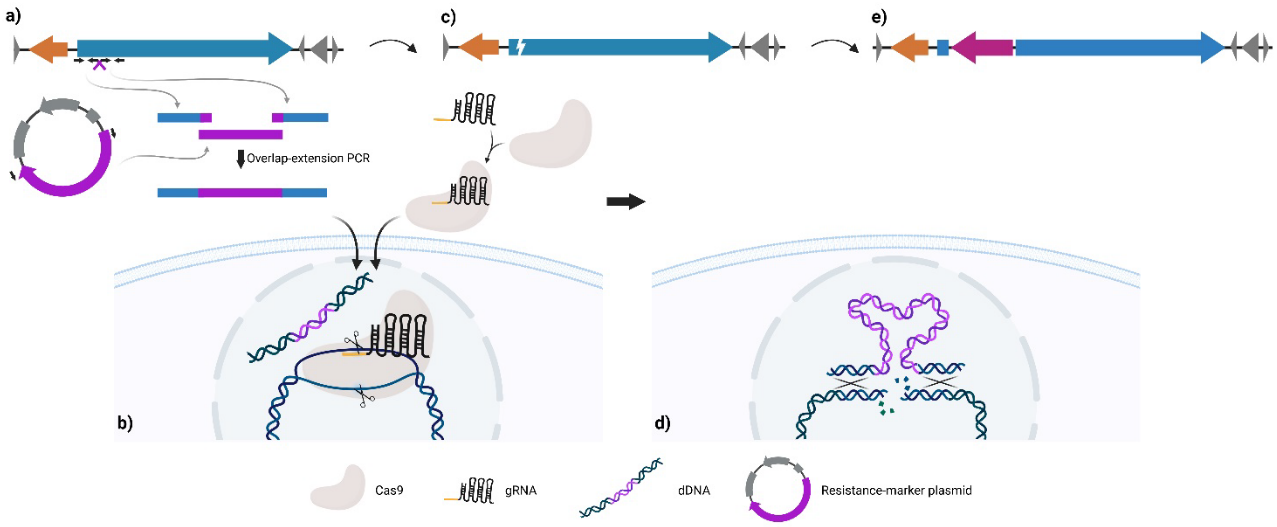

Fig. S15: Schematic of CRISPR/Cas9 genome editing approach in Apiospora.arundinis AAU773. a) A donor DNA (dDNA) cassette is constructed by overlap-extension-PCR of 1.1 kb homology regions of the target exon 1 cut-site and an insertion fragment including a HygR resistance marker. b) The dDNA cassette and in vitro assembled RNP complexes of crRNA::tracrRNA and Cas9 are transformed into protoplasts of the fungus by PEG-mediated transformation. c) The RNP complex introduces a double-stranded break at the target site in the genome. d) the insertion fragment of the dDNA cassette is integrated into the genome at the target cut-site by homology-directed repair facilitated by the cut-site homology bands of the dDNA cassette. e) The nrps0 gene is inactivated by stable integration of the selection marker into the genome (knock-out by knock-in).

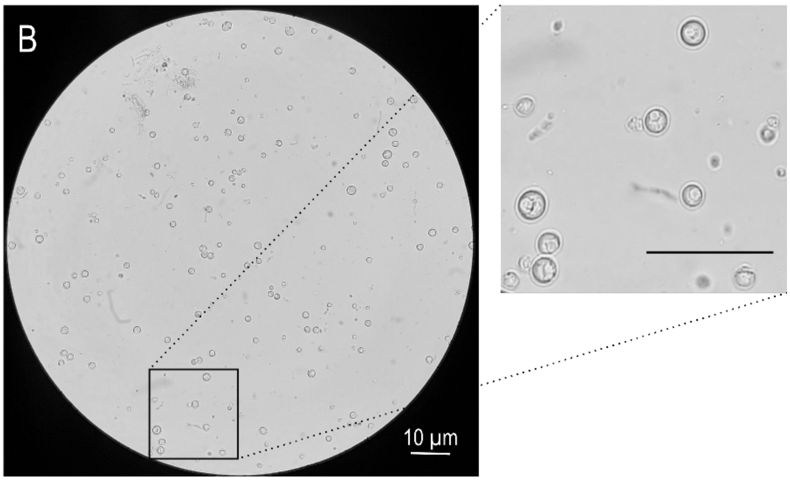

Fig. S16: Protoplastation of 5-day-old Ap¡.arundinis AAU773 on YPG following vortex bead beating homogenization. Digested with 25 mg/mL Glucanex, 65 mg/mL VinoTaste (5) for 3 hr before reaction termination and protoplast purification. Bar = 10 µm.

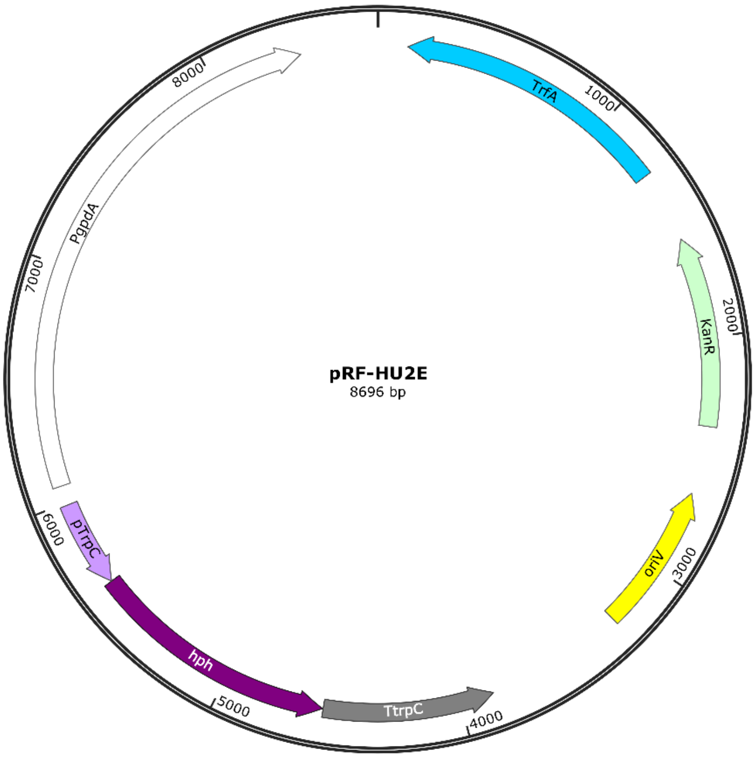

Fig. S17: Plasmid map of the pRF-HU2E vector [1].

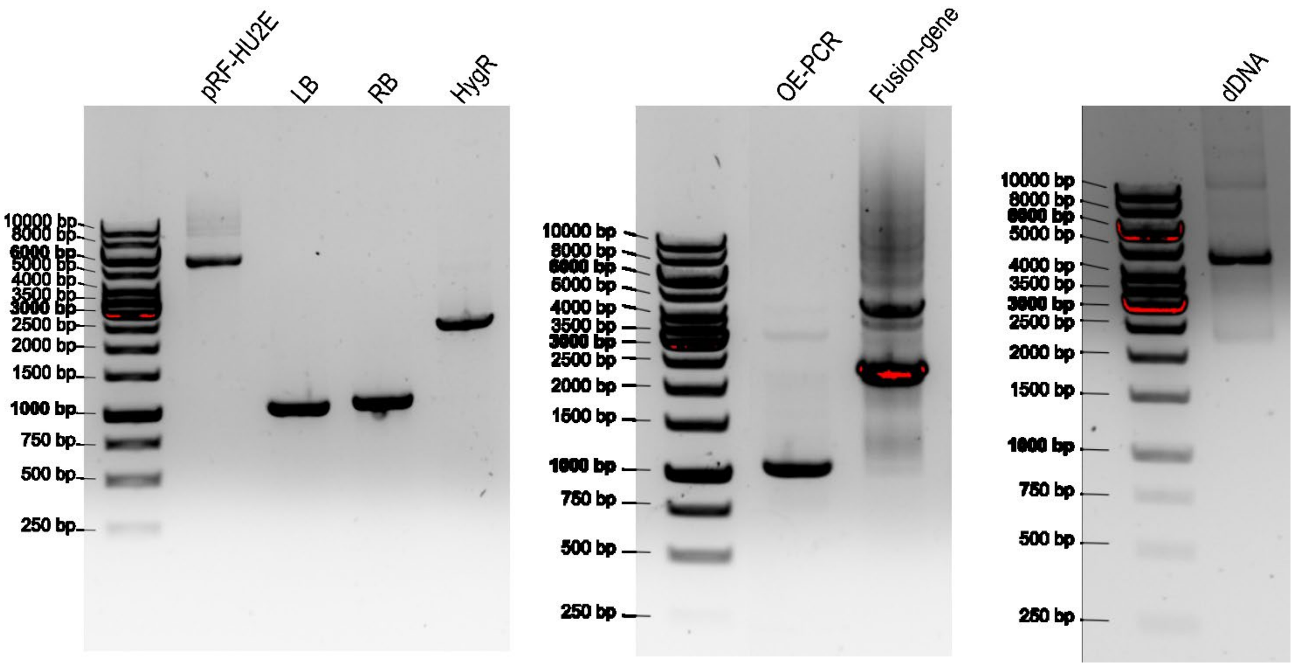

Fig. S18: Agarose gels (1 %) of dDNA constituent fragments and final cassette for *Ap. arundinis* AAU773 Δ*nrps4* mutant generation. Left: purified pRF-HU2E plasmid DNA, left homology band (LB), right homology band (RB), amplified HygR gene (HygR). Middle: overlap extension PCR reaction (used as template for fusion-gene amplification) (OE-PCR), fusion-gene amplification products (fusion-gene). Right: Purified dDNA template following gel extraction (dDNA).

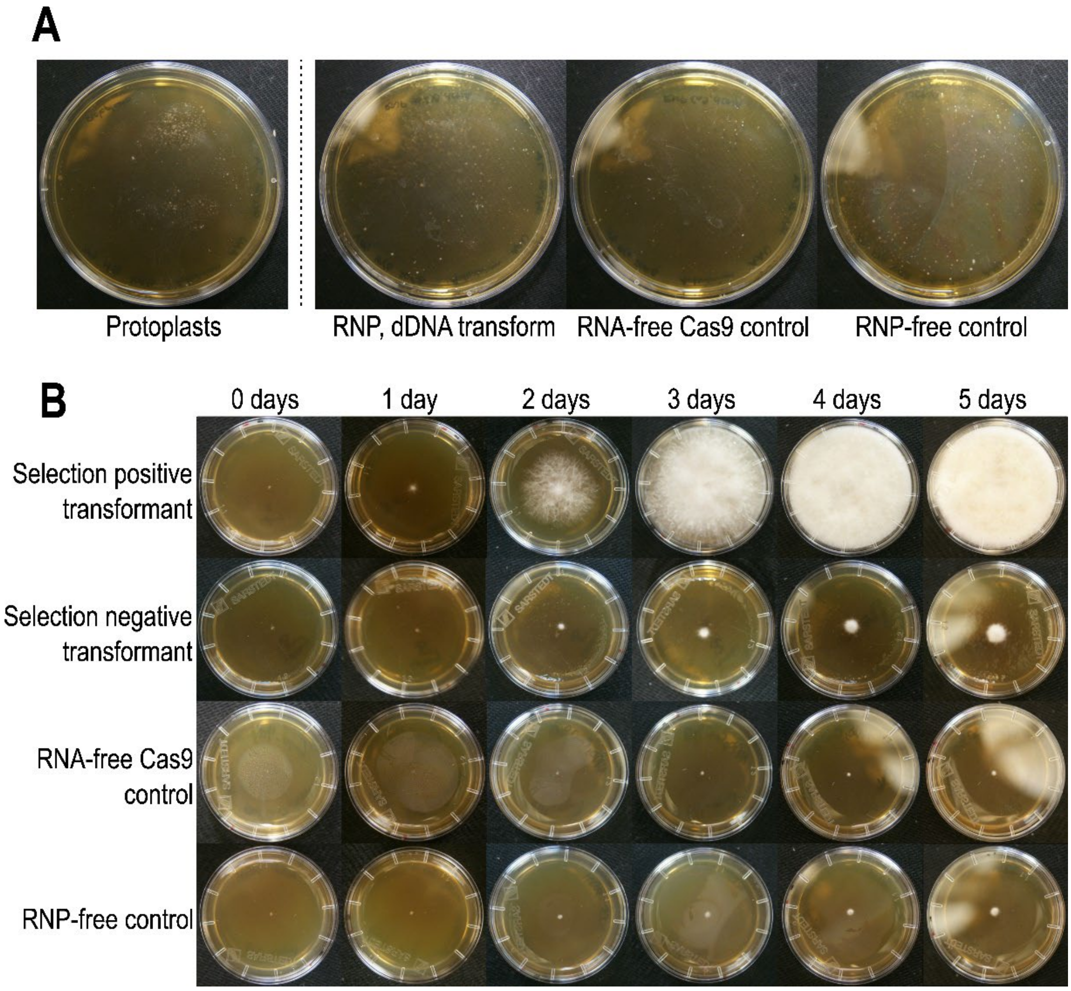

Fig. S19: A) Ap¡.arundinis transformation recovery plates three days after inoculation. B) Colonies extracted from transformation plates on selection YPG medium.

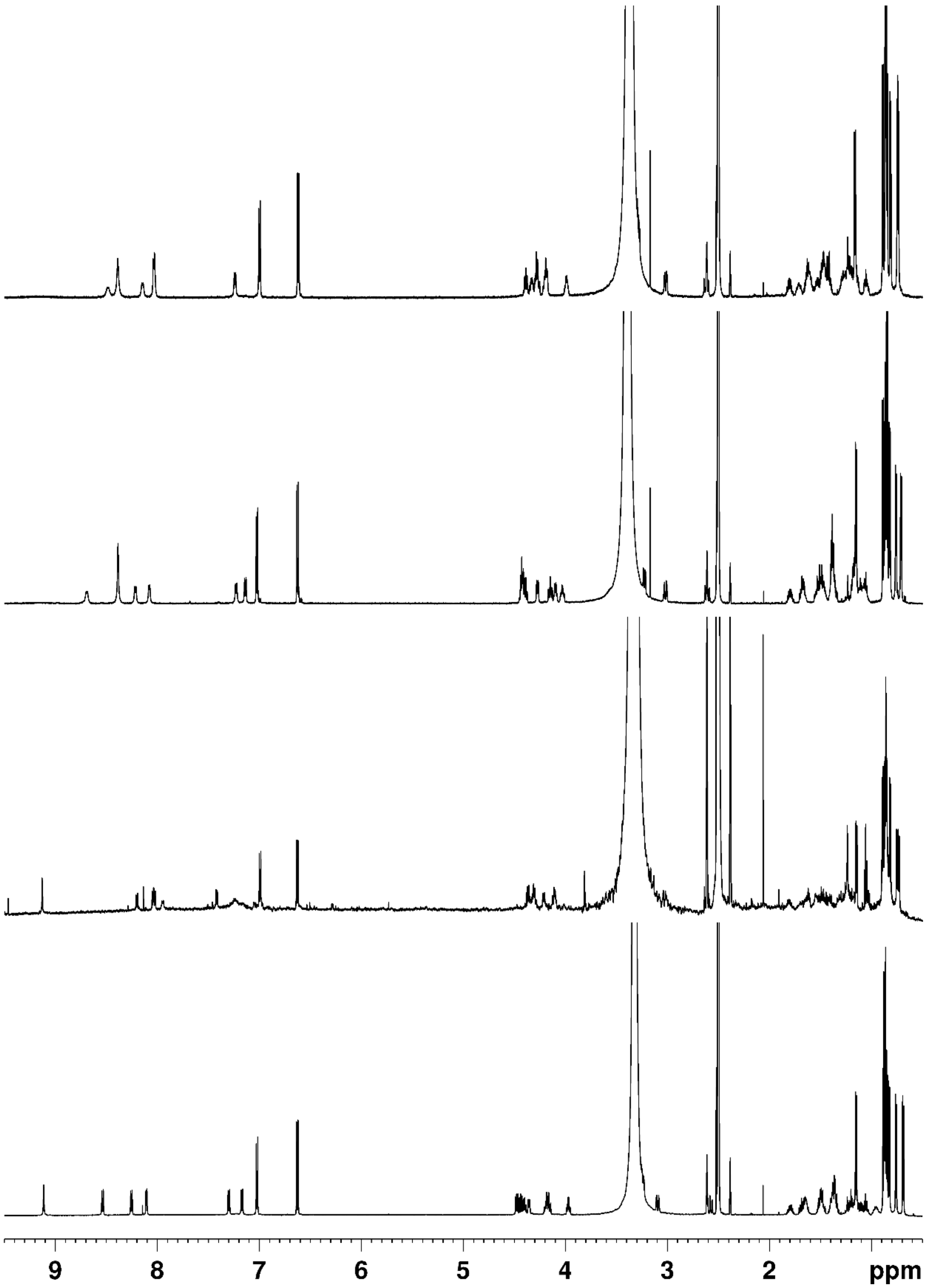

Fig S20. $^{1}$H-NMR spectra of Apiosporin A1-A4 (top to bottom) in DMSO-$d_6$ at 308.1 K.

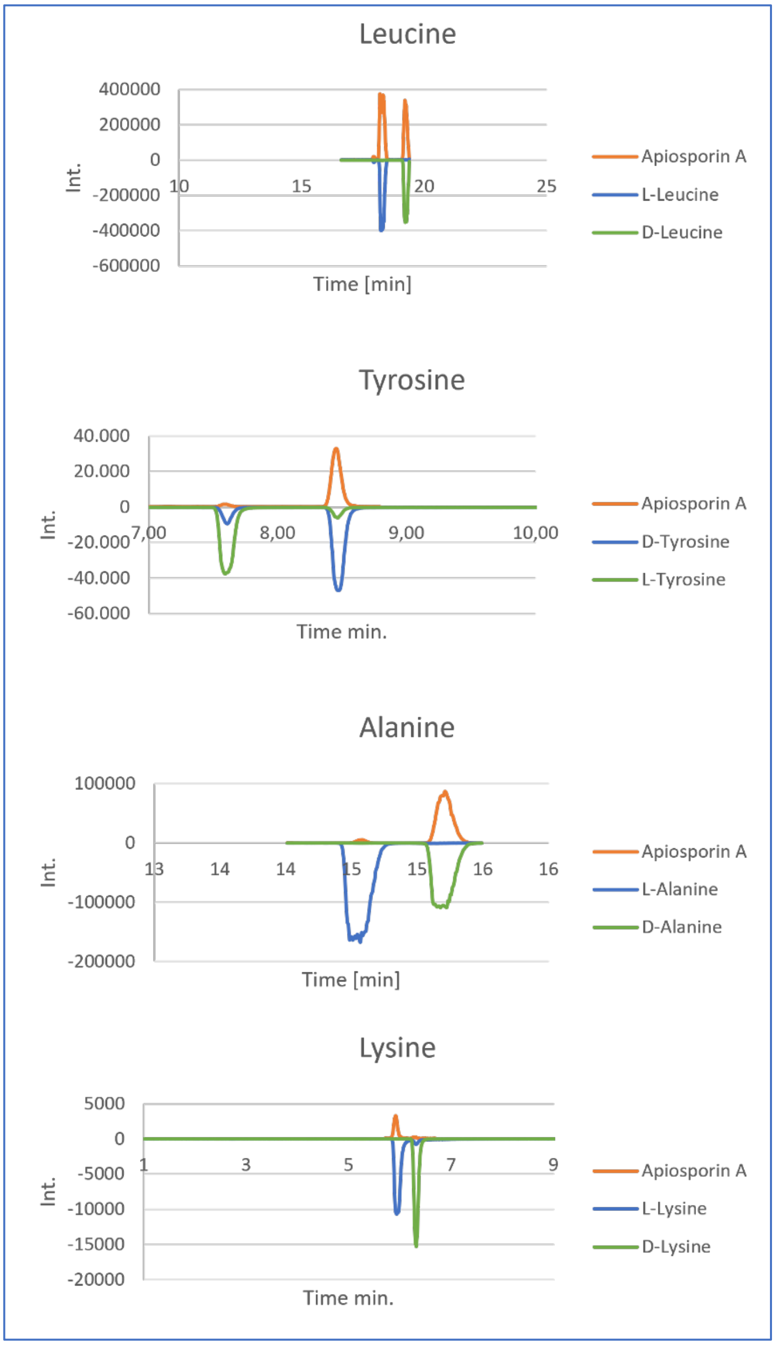

Fig. S21: Marfeys analysis of Apiosporin A. RP-HPLC-HRMS EICs of FDAA derivatized apiosporin A hydrolysate and FDAA derivatized standard amino acids. Lysine was used as a standard for the lysine residue of apiosporin A that has undergone heterocyclization through its side chain amine.

### Supplementary Tables

Tab. S1. WARP biosynthetic gene cluster gene entries. Species and corresponding gene identifiers for WARP BGCs depicted in Fig. S1.

| Class | Species | Strain | Genome | Gene ID |
|---|---|---|---|---|
| Sordariomycetes | Apiospora arundinis | AAU773 | ASM3910530v1 | PGQ11_012562 |
| Sordariomycetes | Chaetomium globosum | CBS 148.51 | ASM14336v1 | CHGG_06789 |
| Sordariomycetes | Emericellopsis atlantica | TS7 | AcreTS7_1 | F5Z01DRAFT_689818 |
| Sordariomycetes | Fusarium graminearum | PH-1; NRRL 31084 | ASM24013v3 | FGSG_02315 |
| Sordariomycetes | Fusarium venenatum | A3/5 | ASM90000737v1 | FVRRES_02708 |
| Sordariomycetes | Metarhizium anisopliae | JEF-290 | ASM1330549v1 | E5D57_001501 |
| Sordariomycetes | Thermothelomyces thermophilus | ATCC 42464 | ASM22609v1 | MYCTH_75351 |
| Eurotiomycetes | Aspergillus fumigatus | Af293 | ASM265v1 | AFUA_1G10380 |
| Eurotiomycetes | Aspergillus nidulans | FGSC A4 | ASM1142v1 | ANIA_00016 |
| Eurotiomycetes | Aspergillus niger | CBS 101883 | Asplac1 | BO96DRAFT_498712 |
| Eurotiomycetes | Trichophyton equinum | CBS 127.97 | ASM15117v1 | TEQG_06632 |
| Eurotiomycetes | Penicillium chrysogenum | IBT35668 | ASM2882703v1 | N7489_004157 |
| Eurotiomycetes | Talaromyces marneffei | 11CN-20-091 | ASM955685v1 | EYB26_007693 |
| Eurotiomycetes | Uncinocarpus reesii | 1704 | ASM351v2 | UREG_04677 |
| Dothideomycetes | Alternaria alternata | SRC1lrK2f | Altal1 | CC77DRAFT_1064818 |
| Dothideomycetes | Cochliobolus heterostrophus (Bipolaris maydis) | ATCC 48331 | CocheC4_1 | PSV08DRAFT_375780 |
| Dothideomycetes | Pseudovirgaria hyperparasitica | CBS 121739 | Psehy1 | EJ05DRAFT_60348 |
| Dothideomycetes | Pyrenophora tritici-repentis | M4 | CUR_PTRM4_2.2 | PtrM4_023140 |
| Dothideomycetes | Trematosphaeria pertusa | CBS 122368 | Trepe1 | BU26DRAFT_424697 |
| Leotiomycetes | Cadophora gregata | MNR1 | UMN_CgA_1 | ONS95_003596 |
| Leotiomycetes | Chlorociboria aeruginascens | IHIA39 | Chloro_aeru_v2 | B7494_g5755 |
| Leotiomycetes | Lachnellula hyalina | CBS 185.66 | CFIA_Lhya_EG2017 | LHYA1_G004496 |
| Leotiomycetes | Venustampulla echinocandica | BP 5553 | ASM335714v1 | BP5553_00199 |
| Leotiomycetes | Glarea lozoyensis | ATCC 20868 | GLAREA | GLAREA_05009 |
| Lecanoromycetes | Bacidia gigantensis | McMullin 20006 | ASM1945646v1 | KY384_005632 |
| Lecanoromycetes | Ramalina farinacea | LIQ254RAFAR | ASM2994837v1 | OHK93_002436 |

Tab. S2. Apiosporin masses and molecular formula. Suggested molecular formula of Ap¡.arundinis. AAU773 NRPS4-derived apiosporin products from $[M+H]^+$ monoisotopic masses.

| **Heptapeptides** | | | | |
|---|---|---|---|---|
| **$[M+H]^+$ [Da]** | **Retention time [min]** | **Molecular formula** | **Δm [ppm]** | **Name** |
| 832.5193 | 5.95 | $C_{42}H_{69}N_7O_{10}$ | 1.06 | |
| 832.5193 | 6.43 | $C_{42}H_{69}N_7O_{10}$ | 1.06 | |
| 814.5069 | 6.50 | $C_{42}H_{67}N_7O_9$ | -1.17 | Apiosporin A |
| 814.5069 | 6.79 | $C_{42}H_{67}N_7O_9$ | -1.17 | Apiosporin A |

| | | | | |
|---|---|---|---|---|
| 874.5664 | 6.63 | $C_{45}H_{75}N_7O_{10}$ | 1.18 | |
| 856.5544 | 7.12 | $C_{45}H_{73}N_7O_9$ | -0.47 | Apiosporin B |
| 856.5544 | 7.43 | $C_{45}H_{73}N_7O_9$ | -0.47 | Apiosporin B |
| 798.5110 | 7.12 | $C_{42}H_{67}N_7O_8$ | -2.43 | |
| 798.5110 | 7.45 | $C_{42}H_{67}N_7O_8$ | -2.43 | |
| 842.5029 | 7.55 | $C_{43}H_{67}N_7O_{10}$ | 0.16 | |
| 840.5588 | 7.74 | $C_{45}H_{73}N_7O_8$ | -1.29 | |
| 840.5588 | 8.10 | $C_{45}H_{73}N_7O_8$ | -1.29 | |
| 884.5510 | 8.28 | $C_{46}H_{73}N_7O_{10}$ | 1.45 | |
| 826.5093 | 8.32 | $C_{43}H_{67}N_7O_9$ | 1.75 | |
| 868.5528 | 9.07 | $C_{46}H_{73}N_7O_9$ | -2.30 | |
| **Hexapeptides** | | | | |
| **[M+H]$^+$ [Da]** | **Retention time [min]** | **Molecular formula** | **Δm [ppm]** | **Name** |
| 687.4447 | 5.98 | $C_{36}H_{58}N_6O_7$ | 0.26 | |
| 705.4549 | 6.00 | $C_{36}H_{60}N_6O_8$ | -0.27 | |
| 687.4447 | 6.27 | $C_{36}H_{58}N_6O_7$ | 0.26 | |
| 687.4447 | 3.52 | $C_{36}H_{58}N_6O_7$ | 0.26 | |
| 747.5018 | 6.53 | $C_{39}H_{66}N_6O_8$ | -0.32 | |
| 687.4447 | 6.76 | $C_{36}H_{58}N_6O_7$ | 0.26 | |
| 729.4914 | 7.15 | $C_{39}H_{64}N_6O_7$ | -0.10 | |
| 729.4914 | 7.41 | $C_{39}H_{64}N_6O_7$ | -0.10 | |

Tab. S3. Proposed apiosporin chemical structures. Proposed structures (without stereochemistry) of Ap¡.arundinis.AAU773.apiosporin products from [M+H]$^+$ monoisotopic masses.

| [M+H]$^+$ [Da] | Proposed compound structure |
|---|---|
| 832.5193 | 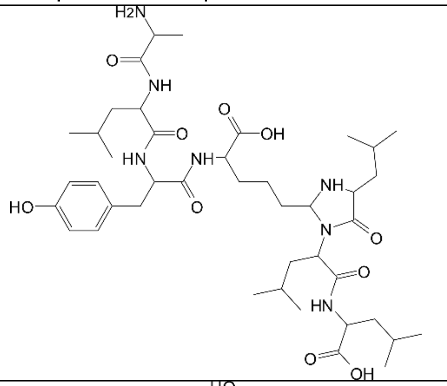 |
| 814.5069 (Apiosporin A) | 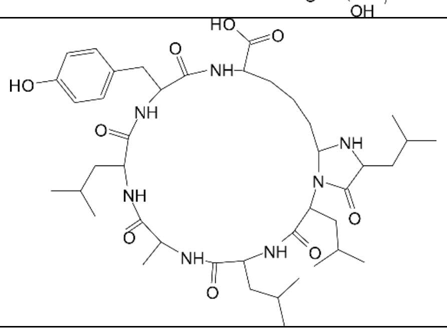 |

| | |
|---|---|
| 874.5664 | 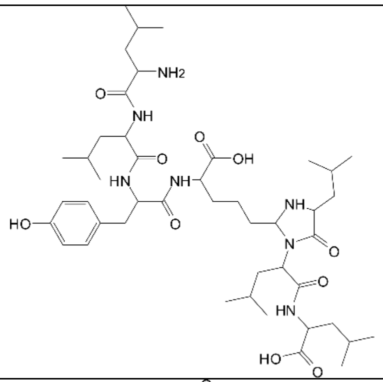 |
| 856.5544 (Apiosporin B) | 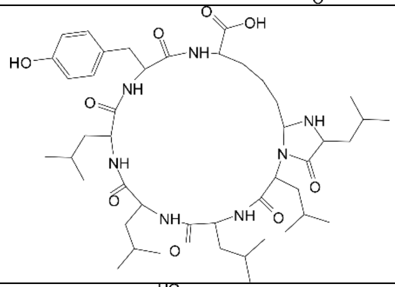 |
| 798.5110 | 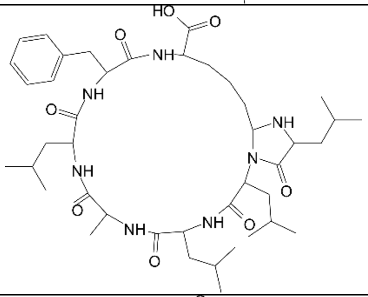 |
| 842.5029 | 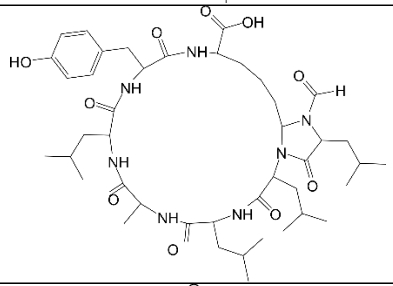 |
| 840.5588 | 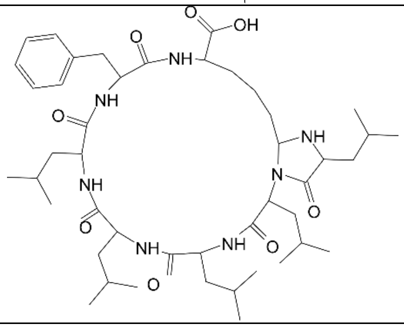 |

| | |
|---|---|
| 884.5510 | 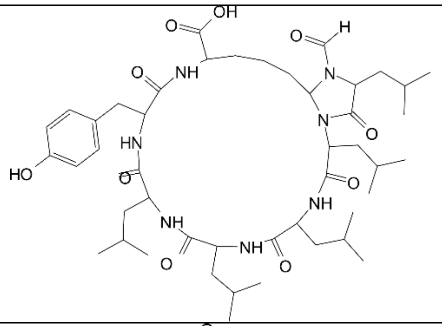 |
| 826.5093 | 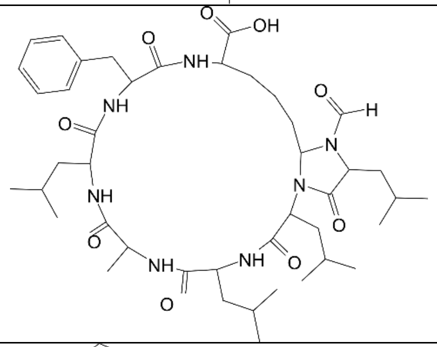 |
| 868.5528 | 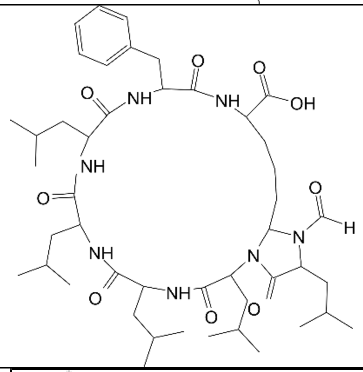 |
| 705.4549 | 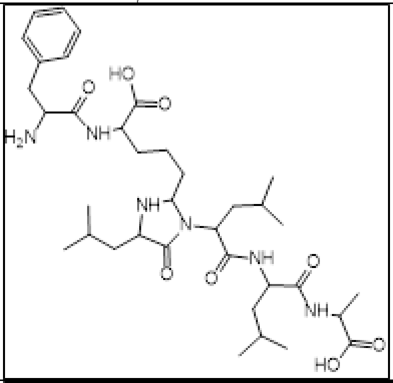 |
| 687.4447 | 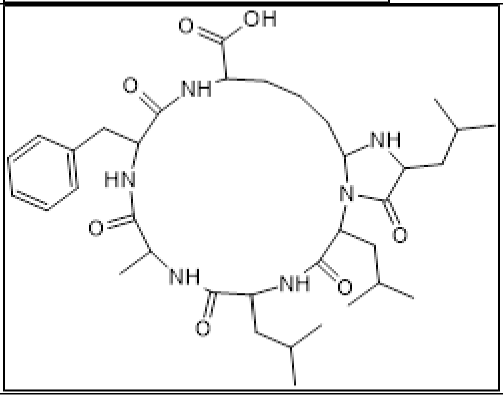 |

| | |
|---|---|
| 747.5018 | 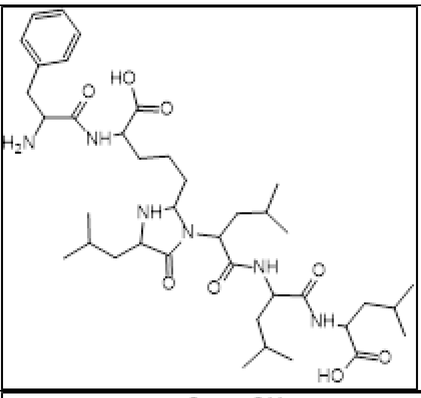 |
| 729.4914 | 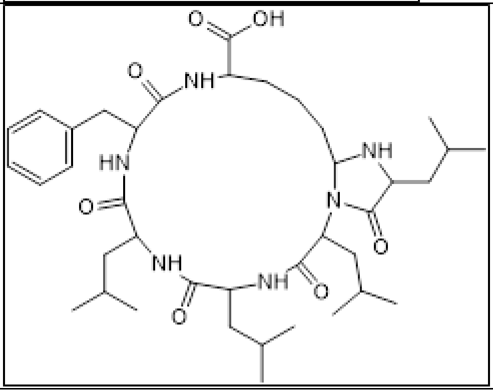 |

**Tab. S4. Confirmed and putative WARP products and producing species.** Gene names are shown for WARPs linked to synthetase genes. *Expected stereochemistry (not confirmed).

| Species | *NRPS* gene | Compound name | Compound amino acid sequence | References |
|---|---|---|---|---|
| **WARP class A – tetrapeptides** | | | | |
| *Aspergillus nidulans*<br>*Aspergillus niger*<br>*Aspergillus versicolor*<br>*Penicillium spp.* | *nlsA*<br>An08g02310<br>-<br>*hcpA* | Fungisporin A | cyclo(Phe(D)-Phe-Val(D)-Val) | [2–9] |
| *Aspergillus nidulans*<br>*Aspergillus niger*<br>*Penicillium spp.* | *nlsA*<br>An08g02310<br>*hcpA* | Fungisporin B | cyclo(Tyr(D)-Phe-Val(D)-Val) | [2,3,5–8,10] |
| *Aspergillus nidulans*<br>*Aspergillus niger*<br>*Aspergillus fumigatus* | *nlsA*<br>An08g02310<br>*pes1* | Fungisporin C | cyclo(Tyr(D)-Tyr-Val(D)-Val)*<br>**cyclo(Tyr(D)-Tyr-Val(D)-Val)*** | [2,3,8]<br>**This study** |

| | | | | |
|---|---|---|---|---|
| *Penicillium verrucosum* | | | | |
| *Aspergillus nidulans* *Aspergillus niger* *Penicillium spp.* | *nlsA* An08g02310 *hcpA* | Fungisporin D | cyclo(Phe(D)-Trp-Val(D)-Val)* | [2,3,5,7,8,10] |
| *Aspergillus niger* *Penicillium chrysogenum* *Penicillium verrucosum* | An08g02310 *hcpA* | | cyclo(Tyr(D)-Trp-Val(D)-Val)* | [3,8] |
| *Aspergillus niger* *Penicillium chrysogenum* | An08g02310 *hcpA* | | cyclo(Phe(D)-Phe-Val(D)-Ile)* | [3] |
| *Aspergillus niger* *Penicillium chrysogenum* | An08g02310 *hcpA* | | cyclo(Phe(D)-Phe-Ile(D)-Val)* | [3] |
| *Aspergillus niger* *Penicillium chrysogenum* *Penicillium rubens* | An08g02310 *hcpA* | | cyclo(Tyr(D)-Trp-Val(D)-Ile)* | [3,5] |
| *Aspergillus niger* *Penicillium chrysogenum* | An08g02310 *hcpA* | | cyclo(Tyr(D)-Trp-Ile(D)-Val)* | [3] |
| *Aspergillus niger* *Penicillium chrysogenum* | An08g02310 *hcpA* | | cyclo(Tyr(D)-Phe-Val(D)-Ile)* | [3] |
| *Aspergillus niger* *Penicillium chrysogenum* | An08g02310 *hcpA* | | cyclo(Tyr(D)-Phe-Ile(D)-Val)* | [3] |
| **WARP class B – pentapeptides** | | | | |
| *Moltirella alpina* | | Malpibaldin A | cyclo(Val(D)-Leu-Leu(D)-Phe(D)-Leu) | [11,12] |
| *Moltirella alpina* | | Malpibaldin B | cyclo(Val(D)-Leu-Leu(D)-Trp(D)-Leu) | [11,12] |
| *Moltirella alpina* | | Malpibaldin C | cyclo(Val-Leu-Leu-Tyr-Leu) | [12] |

| | | | | |
|---|---|---|---|---|
| **WARP class C – hexapeptides** | | | | |
| *Fusarium graminearum* | *nrps4* | Fusahexin | cyclo(Ala(D)-Leu-*allo*-Thr(D)-Pro-Leu(D)-Leu) | [13] |
| *Fusarium solani* | | | cyclo(Hyp-Xle-Xle-Ala-Thr-Xle) cyclo(Dhp-Xle-Xle-Ala-Thr-Xle) cyclo(Hyp-Xle-Xle-Val-Thr-Xle) cyclo(Dhp-Xle-Xle-Val-Thr-Xle) | [14] |
| *Fusarium roseum Fusarium tricinctum* | | Acuminatum A | cyclo(3S,4R-HMTA-allo-Thr(D)-Ala-Ala(D)-Gln-Tyr(D)-Leu) | [15,16] |
| *Fusarium roseum Fusarium tricinctum Fusarium spp.* | | Acuminatum B | cyclo(3S,4R-HMTA-allo-Thr(D)-Ala-Ala(D)-Gln-Tyr(D)-Ile) | [15–17] |
| *Fusarium roseum Fusarium tricinctum Fusarium spp.* | | Acuminatum C | cyclo(3S,4R-HMTA-allo-Thr(D)-Ala-Ala(D)-Gln-Tyr(D)-Val) | [15–17] |
| **WARP class D – heptapeptides** | | | | |
| *Apiospora arundinis* | PGQ11_012562 *nrps4* | Apiosporin A | **cyclo(**Leu-Leu(D)-Leu-Ala(D)-Leu-Tyr(D)-Lys**)** | **This study** |
| *Apiospora arundinis* | PGQ11_012562 *nrps4* | Apiosporin B | **cyclo(**Leu-Leu(D)-Leu-Leu(D)-Leu-Tyr(D)-Lys**)** | **This study** |
| **WARP class E – octapeptides** | | | | |

Tab. S5: List of primers (As¡.fumigatus).

| Name | Annealing region (5'-3') (5' overhang in red) |
|---|---|
| Primer 1 | TCCTTAAGAAGAAACCTGCTTGGGAGTACTTTCAGGTACTTCGACCCCTCTCGTGGACCTAGCTGATTCT |
| Primer 2 | CTCTCCCGACAGTCCTCTCGAGATAGACAGAAGCTCACGCAAAGCCTCTGATCATCATGCAACATGCATG |
| Primer 3 | GGCGAGGTAGGGTGGCTCAGAA |
| Primer 4 | TTCTGCGGGCGATTTGTG |

| | |
|---|---|
| Primer 5 | AACTTTGGTGCTACCTGTGATG |
| Primer 6 | GAGGCGATGTTCGGGGAT |

Tab. S6: Protospacer sequence and metadata (As¡.fumigatus).

| Name | Sequence (5’-3’) | PAM | Strand |
|---|---|---|---|
| Protospacer 1 | GTAAAGCCGAGCATTTTCAG | AGG | (+) |
| Protospacer 2 | AGGCATCACCACAACCCGAG | GGG | (-) |

Tab. S7: Assembly statistics of As¡.fumigatus.⊥pes7.

| Strain | Contigs | Genome size (Mb) | N50 (Mb) | N99 (Mb) | Reference |
|---|---|---|---|---|---|
| As¡.fumigatus.⊥pes7 | 23 | 29.03 | 1 | 0 | This study |
| As¡.fumigatus | 10 | 28.96 | 3 | | This study |

Tab. S8: $^{1}$H and $^{13}$C-chemical shifts of cyclo(Tyr(D)-Tyr-Val(D)-Val) in DMSO-$d_6$ at 308.1 K:

| | | | | | | | | | |
|---|---|---|---|---|---|---|---|---|---|
| **Tyr (D) 1** | $H^N$ | 7.98 | C’ | 172.53 | **Val(D) 3** | $H^N$ | 7.72 | C’ | 172.95 |
| | $H^\alpha$ | 4.44 | $C^\alpha$ | 53.00 | | $H^\alpha$ | 3.89 | $C^\alpha$ | 58.73 |
| | $H^{\beta 2}$ | 2.63 | $C^\beta$ | 33.34 | | $H^\beta$ | 1.92 | $C^\beta$ | 26.49 |
| | $H^{\beta 3}$ | 2.83 | $C^\gamma$ | 127.71 | | $H^{\gamma 1}$ | 0.78 | $C^{\gamma 1}$ | 18.87 |
| | $H^\delta$ | 6.96 | $C^\delta$ | 129.49 | | $H^{\gamma 2}$ | 0.73 | $C^{\gamma 2}$ | 18.15 |
| | $H^\varepsilon$ | 6.59 | $C^\varepsilon$ | 114.61 | | | | | |
| | | | $C^\zeta$ | 155.53 | | | | | |
| **Tyr 2** | $H^N$ | 7.74 | C’ | 172.97 | **Val 4** | $H^N$ | 7.59 | C’ | 172.66 |
| | $H^\alpha$ | 4.46 | $C^\alpha$ | 53.41 | | $H^\alpha$ | 3.89 | $C^\alpha$ | 58.73 |
| | $H^{\beta 2}$ | 2.59 | $C^\beta$ | 33.89 | | $H^\beta$ | 1.90 | $C^\beta$ | 26.66 |
| | $H^{\beta 3}$ | 2.87 | $C^\gamma$ | 127.68 | | $H^{\gamma 1}$ | 0.81 | $C^{\gamma 1}$ | 18.18 |
| | $H^\delta$ | 6.94 | $C^\delta$ | 129.41 | | $H^{\gamma 2}$ | 0.70 | $C^{\gamma 2}$ | 18.75 |
| | $H^\varepsilon$ | 6.59 | $C^\varepsilon$ | 114.61 | | | | | |
| | | | $C^\zeta$ | 155.52 | | | | | |

Tab. S9: List of primers (Ap¡.arundinis)

| Name | Annealing region (5’-3’) |
|---|---|

| | (5’ overhang in red) |
|---|---|
| Primer 1 | CCATGCATACACGTACATGCCTAGTATCAT |
| Primer 2 | TGGCGCCTATATCGCCGACATCACCGATCATTACTACGCATCACATCTTTCACCAGCACACGTATG |
| Primer 3 | TATCTCTACACACAGGCTCAAATCAATTAGTAATGAAATTCAGGGGCTCGCATCAGCTGCCAAGCA |
| Primer 4 | TGATGGACTCAATGGAGTCGATTGGTGCAT |
| Primer 5 | TAGTAATGATCGGTGATGTCGGCGATATA |
| Primer 6 | TCATTACTAATTGATTTGAGCCTGTGTGT |

Tab. S10: Protospacer sequence and metadata (Ap¡.arundinis).

| Name | Sequence (5’-3’) | PAM | Strand | On-target score |
|---|---|---|---|---|
| Protospacer 1 | CTGGTGAAAGATGTGATGCG | AGG | (+) | 68.8 |

Tab. S11: Assembly statistics of Ap¡.arundinis.AAU773 Δnrps0

| Strain | Contigs | Genome size (Mb) | N50 (Mb) | N99 (Mb) | Reference |
|---|---|---|---|---|---|
| Ap¡.arundinis.AAU773 Δnrps0 | 15 | 48.94 | 5.30 | 0.75 | This study |
| Ap¡.arundinis.AAU773 | 10 | 48.8 | 5.3 | 3.0 | [18] |

Tab. S12a: $^{1}$H, $^{13}$C and $^{15}$N-chemical shifts of Apiosporin A1-A4, in DMSO-$d_6$ at 308.1 K. Spectra were calibrated against external TMS (0.00 ppm for $^{1}$H and $^{13}$C. Shifts of $^{15}$N were calibrated against $NH_3$ = 0.00 ppm[19]. The concentrations of the four samples were determined by using the residual intensity of DMSO (% D) = 98.85% mM as internal standard. A1: 8.5 mM, A2 : 9.2 mM, A3 : 0.7 mM, A4 : 5.3 mM.

| Leu 1 | Position | A1 | A2 | A3 | A4 |
|---|---|---|---|---|---|
| | N | 47.95 | 47.44 | | |
| | C | 175.14 | 176.86 | 174.92 | 176.47 |
| | Cα | 55.74 | 56.80 | 55.64 | 56.62 |
| | Cβ | 40.59 | 41.2 | 40.48 | 41.17 |
| | Cγ | 24.52 | 24.71 | 24.52 | 24.69 |
| | Cδ1 | 23.19 | 23.24 | 23.14 | 23.22 |
| | Cδ2 | 21.68 | 21.26 | 21.62 | 21.38 |
| | Hα | 3.28 | 3.22 | 3.27 | 3.24 |
| | Hβ2 | 1.48 | 1.51 | 1.49 | 1.50 |
| | Hβ3 | 1.22 | 1.36 | 1.22 | 1.36 |
| | Hγ | 1.80 | 1.80 | 1.81 | 1.79 |
| | Hδ1 | 0.89 | 0.89 | 0.89 | 0.88 |
| | Hδ2 | 0.86 | 0.86 | 0.86 | 0.85 |
| Leu 2 | N | 137.42 | 136.17 | n.d. | 136.42 |
| | C | 169.12 | 169.95 | 168.99 | 169.72 |

| | | | | | |
|---|---|---|---|---|---|
| | Cα | 54.42 | 52.20 | 54.10 | 51.86 |
| | Cβ | 38.93 | 35.86 | 38.99 | 35.55 |
| | Cγ | 24.68 | 24.47 | 24.71 | 24.46 |
| | Cδ1 | 22.30 | 22.72 | 22.28 | 22.52 |
| | Cδ2 | 22.30 | 21.73 | 22.22 | 21.83 |
| | Hα | 4.29 | 4.42 | 4.31 | 4.47 |
| | Hβ2 | 1.63 | 1.68 | 1.61 | 1.68 |
| | Hβ3 | 1.60 | 1.53 | 1.61 | 1.47 |
| | Hγ | 1.42 | 1.38 | 1.43 | 1.38 |
| | Hδ1 | 0.86 | 0.87 | 0.86 | 0.87 |
| | Hδ2 | 0.85 | 0.82 | 0.85 | 0.82 |
| Leu3 | N | 122.45 | 118.93 | 117,96 | 118.97 |
| | C | 170.98 | 170.44 | 171.58 | 170.56 |
| | Cα | 50.80 | 50.53 | 50.58 | 50.36 |
| | Cβ | 40.04 | 41.82 | 40.05 | 42.23 |
| | Cγ | 24.19 | 24.26 | 24.15 | 24.46 |
| | Cδ1 | 22.95 | 23.08 | 22.94 | 21.65 |
| | Cδ2 | 21.37 | 21.70 | 21.21 | 23.16 |
| | HN | 8.039 | 7.22 | 8.100 | 7.169 |
| | Hα | 4.34 | 4.39 | 4.35 | 4.40 |
| | Hβ2 | 1.46 | 1.38 | 1.47 | 1.38 |
| | Hβ3 | 1.43 | 1.38 | 1.40 | 1.34 |
| | Hγ | 1.54 | 1.48 | 1.54 | 1.51 |
| | Hδ1 | 0.86 | 0.84 | 0.86 | 0.86 |
| | Hδ2 | 0.81 | 0.85 | 0.81 | 0.85 |
| Ala 4 | N | 121.25 | 121.77 | 120.76 | 121.5 |
| | C | 171.85 | 172.13 | 171.91 | 172.50 |
| | Cα | 47.36 | 47.18 | 47.34 | 47.05 |
| | Cβ | 19.12 | 19.56 | 19.18 | 19.50 |
| | HN | 8.15 | 8.22 | 8.02 | 8.26 |
| | Hα | 4.39 | 4.43 | 4.38 | 4.43 |
| | Hβ | 1.16 | 1.15 | 1.14 | 1.15 |
| Leu 5 | N | 118.74 | 119.49 | 122,23 | 119.89 |
| | C | 171.64 | 171.99 | 171.58 | 172.13 |
| | Cα | 51.12 | 51.70 | 51.31 | 52.21 |
| | Cβ | 40.83 | 40.56 | 40.75 | 40.20 |
| | Cγ | 23.94 | 23.85 | 23.78 | 23.64 |
| | Cδ1 | 21.67 | 21.99 | 22.61 | 22.34 |
| | Cδ2 | 22.70 | 22.51 | 21.76 | 22.23 |
| | H | 8.05 | 8.06 | 8.10 | 8.12 |
| | Hα | 4.21 | 4.10 | 4.103 | 3.97 |
| | Hβ2 | 1.15 | 1.11 | 1.17 | 1.20 |
| | Hβ3 | 1.05 | 1.18 | 1.05 | 1.15 |

| | | | | | |
|---|---|---|---|---|---|
| | Hγ | 1.23 | 1.17 | 1.184 | 1.10 |
| | Hδ1 | 0.74 | 0.71 | 0.749 | 0.76 |
| | Hδ2 | 0.75 | 0.76 | 0.730 | 0.69 |
| Tyr 6 | N | 118.67 | 119.80 | 118.06 | 119.68 |
| | C | 170.30 | 170.38 | 171.16 | 170.85 |
| | Cα | 55.46 | 55.90 | 54.88 | 55.42 |
| | Cβ | 36.26 | 35.78 | 36.12 | 35.44 |
| | Cγ | 128.28 | 128.50 | 127.79 | 128.31 |
| | Cδ | 129.85 | 129.80 | 129.78 | 129.74 |
| | Cε | 114.78 | 114.79 | 114.79 | 114.78 |
| | Cζ | 15572 | 155.74 | 155.77 | 155.72 |
| | HN | 8.55 | 8.65 | 8.26 | 8.54 |
| | Hα | 4.27 | 4.15 | 4.301 | 4.17 |
| | Hβ2 | 3.02 | 3.02 | 3.024 | 3.10 |
| | Hβ3 | 2.62 | 2.61 | 2.61 | 2.58 |
| | Hδ | 7.00 | 7.03 | 6.99 | 7.02 |
| | Hε | 6.62 | 6.63 | 6.62 | 6.63 |
| | Hη | 9.15(br) | 9.17(br) | | 9.13 |
| Lys 7 | N | 118.6 | 118.22 | 114.91 | 114.91 |
| | C | 174.74 | 175.08 | n.d. | 173.45 |
| | Cα | 53.97 | 53.01 | 52.49 | 51.24 |
| | Cβ | 32.05 | 31.66 | 30.74 | 30.39 |
| | Cγ | 21.11 | 19.71 | 21.45 | 18.88 |
| | Cδ | 34.51 | 33.75 | 33.99 | 33.09 |
| | Cε | 73.42 | 71.34 | 73.04 | 70.91 |
| | HN | 7.24 | 7.15 | 7.46 | 7.30 |
| | Hα | 3.99 | 4.03 | 4.11 | 4.19 |
| | Hβ2 | 1.71 | 1.67 | 1.55 | 1.64 |
| | Hβ3 | 1.47 | 1.46 | 1.70 | 1.64 |
| | Hγ2 | 1.26 | 1.07 | 1.32 | 1.05 |
| | Hγ3 | 1.18 | 1.07 | 1.26 | 0.96 |
| | Hδ2 | 1.62 | 1.39 | 1.65 | 1.38 |
| | Hδ3 | 1.28 | 1.17 | 1.31 | 1.19 |
| | Hε | 4.18 | 4.28 | 4.21 | 4.35 |

Tab. S12b: Observed homonuclear and heteronuclear couplings and the chemical shifts ($^{1}H$ and $^{13}C$: δ(TMS)=0.00 ppm, $^{15}N$: δ($NH_3$)=0.00 ppm) of Apiosporin A1 in DMSO-$d_6$, at 308 K.

| position | δ ($^{13}C$) [ppm] | m ($^{13}C$) | δ ($^{1}H$) [ppm] | $J_{HH}$ | $^{n}J_{XH}$ |
|---|---|---|---|---|---|
| 1-Leu CO | 175,14 | C | | | |

| | | | | | |
|---|---|---|---|---|---|
| 1-Leu-α | 55,74 | CH | 3,281 | 1,48 (β1), 1,21 (β2) | 175,14 (CO), 169,15 (2-Leu-CO), 47.95 (1-Leu-N) |
| 1-Leu-β | 40,59 | $CH_2$ | 1,476<br>1,215 | 1,22 (β2),3,28 (α)<br>1,47 (β1),3,27 (α) | 23,19 (δ1), 21,68 (δ2), 24,52 (γ), 47.95 (1-Leu-N)<br>23,19 (δ1), 21,69 (δ2) |
| 1-Leu-γ | 24,52 | CH | 1,803 | 1,48 (β1), 1,22 (β2), 0,89 (δ1), 0,86 (δ2 | 23,19 (δ1), 21,68 (δ2) |
| 1-Leu-δ1 | 23,19 | $CH_3$ | 0,889 | 1,80 (γ) | 21,68 (δ2), 24,51 (γ) |
| 1-Leu-δ2 | 21,68 | $CH_3$ | 0,862 | 1,89 (γ) | 24,51 (γ), 23,20 (δ1) |
| 1-Leu-N | | N | | | |
| 2-Leu-CO | 169,12 | C | | | |
| 2-Leu-α | 54,42 | CH | 4,289 | 1,63 (β1), 1,59 (β2) | |
| 2-Leu-β | 38,93 | $CH_2$ | 1,634<br>1,597 | 4,29 (α)<br>4,29 (α) | 169,12 (CO), 22,3 (δ1,2), 137,42 (2-Leu-N)<br>169,12 (CO), 22,3 (δ1,2), 24,68 (γ), |
| 2-Leu-γ | 24,68 | CH | 1,424 | 0,86 (δ1), 0,84 (δ2) | 22,29 (δ1,2) |
| 2-Leu-δ1 | 22,30 | $CH_3$ | 0,859 | 1,42 (γ) | 22,30 (δ2), 24,68 (γ) |
| 2-Leu-δ2 | 22,30 | $CH_3$ | 0,846 | 1,42 (γ) | 24,68 (γ), 22,30 (δ1) |
| 2-Leu-N | | N | | | |
| 3-Leu-CO | 170,98 | C | | | |
| 3-Leu-α | 50,80 | CH | 4,337 | 8,04 (NH), 1,46 (β1), 1,42 (β2) | 170,98 (CO), 169,12 (2-Leu-CO) |
| 3-Leu-β | 40,04 | $CH_2$ | 1,455<br>1,425 | 4,34 (α)<br>4,34 (α) | 24,18 (γ), 170,89 (CO), 21,37 (δ2)<br>21,36 (δ2), 24,1 (γ) |
| 3-Leu-γ | 24,19 | CH | 1,536 | 0,86 (δ1), 0,81 (δ2) | |
| 3-Leu-δ1 | 22,95 | $CH_3$ | 0,857 | 1,54 (γ) | 21,37 (δ2) 24,19 (γ) |
| 3-Leu-δ2 | 21,37 | $CH_3$ | 0,814 | 1,54 (γ) | 22,95 (δ1), 24,18 (γ) |
| 3-Leu-NH | 122,45 | NH | 8,039 | 4,34 (α) | 169,13 (2-Leu-CO) |

| | | | | | |
|---|---|---|---|---|---|
| 4-Ala-CO | 171,85 | C | | | |
| 4-Ala-α | 47,36 | CH | 4,393 | 8,15 (NH), 1,16 (β) | 171,85 (CO), 170,98 (3-Leu-CO) |
| 4-Ala-β | 19,12 | $CH_3$ | 1,163 | 4,39 (α) | 171,83 (CO), 121,25 (4-Ala-N) |
| 4-Ala-NH | 121,25 | NH | 8,154 | 4,39 (α) | 122,45 (3-Leu-N) |
| 5-Leu-CO | 171,64 | C | | | |
| 5-Leu-α | 51,12 | CH | 4,213 | 8,04 (NH), 1,14 (β1), 1,04 (β2) | 171,66 (CO) |
| 5-Leu-β | 40,83 | $CH_2$ | 1,147<br>1,054 | 1,05 (β2), 4,21 (α), 1.23 (γ)<br>1,14 (β1), 4,21 (α), 1,23 (γ) | 21,67 (δ2), 22,68 (δ1), 23,95 (γ)<br>23,92 (γ), 21,67 (δ2), 22,69 (δ1), 118.74 (5-Leu-N) |
| 5-Leu-γ | 23,94 | CH | 1,231 | 1,14 (β1), 1,05 (β2),0,74 (δ1), 0,74 (δ2) | 22,70 (δ1) |
| 5-Leu-δ1 | 22,70 | $CH_3$ | 0,745 | 1,23 (γ) | 21,67 (δ2), 23,93 (γ) |
| 5-Leu-δ2 | 21,67 | $CH_3$ | 0,738 | 1,23 (γ) | 22,68(δ1), 23,93 (γ), |
| 5-Leu-NH | 118,74 | NH | 8,046 | 4,21 (α) | 171,84 (4-Ala-CO) |
| 6-Tyr-CO | 170,30 | C | | | |
| 6-Tyr-α | 55,46 | CH | 4,266 | 8,55 (NH), 3,01 (β1), 2,62 (β2) | 128,28 (γ), 170,30 (CO), 171,64 (5-Leu-CO) |
| 6-Tyr-β | 36,26 | $CH_2$ | 3,017<br>2,621 | 4,27 (α), 2,62 (β2)<br>4,27 (α), 3,02 (β1) | 170,34 (CO), 128,26 (γ), 129,85 (δ), 118.67 (6-Tyr-N)<br>170,34 (CO), 128,26 (γ), 129,85 (δ), |
| 6-Tyr-γ | 128,28 | C | | | |
| 6-Tyr-δ,δ´ | 129,85 | CH | 7,002 | 6,62 (ε) | 129,85 (δ´), 155,73 (ζ) |
| 6-Tyr-ε,ε´ | 114,78 | CH | 6,619 | 7,00 (δ) | 155,72 (ζ), 128,28 (γ), |
| 6-Tyr-ζ | 155,72 | C | | | |
| 6-Tyr-OH | | OH | 9,15 βr | | |
| 6-Tyr-NH | 118,67 | NH | 8,553 | 4,26 (α) | 170,31 (6-Tyr-CO) |
| 7-Lys-CO | 174,74 | C | | | |

| | | | | | |
|---|---|---|---|---|---|
| 7-Lys-α | 53,97 | CH | 3,987 | 7,24 (NH), 1,71 (β1), 1,47 (β2) | 21,12 (γ), 170,30 (6-Tyr-CO), 174,74 (CO) |
| 7-Lys-β | 32,05 | $CH_2$ | 1,712<br>1,473 | 3,98 (α), 1,47 (β2), 1,26 (γ1), 1,18 (γ2)<br>3,98 (α), 1,71 (β2), 1,26 (γ1), 1,18 (γ2) | |
| 7-Lys-γ | 21,11 | $CH_2$ | 1,258<br>1,184 | 1,62 (γ1), 1,71 (β1), 1,47 (β2),1,18 (γ2)<br>1,26 (γ1), 1,71 (β1), 1,47 (β2), 1,62 (δ1) | |
| 7-Lys-δ | 34,51 | $CH_2$ | 1,621<br>1,284 | 1,28 (δ2), 1,18 (γ2), 4,18 (ε)<br>1,62 (δ1), 4,18 (ε) | |
| 7-Lys-ε | 73,42 | CH | 4,175 | 1,62 (δ1), 1,28 (δ2) | 175,16 (1-Leu-CO) |
| 7-Lys-NH | 118,71 | NH | 7,242 | 3,98 (α) | |

Tab. S12c: Observed homonuclear and heteronuclear couplings and the chemical shifts ($^{1}H$ and $^{13}C$: δ(TMS)=0.00 ppm, $^{15}N$: δ($NH_3$)=0.00 ppm) of Apiosporin A2 in DMSO-$d_6$, at 308 K.

| position | δ ($^{13}C$,$^{15}N$) [ppm] | m ($^{13}C$, $^{15}N$) | δ ($^{1}H$) [ppm] | $J_{HH}$ | $^{n}J_{XH}$ |
|---|---|---|---|---|---|
| 1-Leu CO | 176,858 | C | | | |
| 1-Leu-α | 56,800 | CH | 3,222 | 1,51 (β1), 1,36 (β2) | 176,86 (1-Leu-CO), 41,13 (1-Leu-β), 71,34 (7-Lys-e) |
| 1-Leu-β | 41,122 | $CH_2$ | 1,510<br>1,359 | 3,22 (α), 1,36 (β2), 1,80 (γ)<br>3,22 (α(, 1,51 (β1), 1,80 (γ) | 47,44 (1-Leu-N), 176,86 (1-Leu-CO), 56,80 (1-Leu-α)<br>47,44 (1-Leu-N), 176,86 (1-Leu-CO), 56,80 (1-Leu-α) |
| 1-Leu-γ | 24,709 | CH | 1,800 | 1,51 (β1), 1,36 (β2), 0,88 (δ1), 0,86 (δ2) | 56,80 (1-Leu-α)<br>41,12 (1-Leu-β) |
| 1-Leu-δ1 | 23,237 | $CH_3$ | 0,888 | 1,80 (γ) | 41,12 (1-Leu-β) |
| 1-Leu-δ2 | 21,463 | $CH_3$ | 0,860 | 1,80 (γ) | 41,12 (1-Leu-β) |

| | | | | | |
|---|---|---|---|---|---|
| 1-Leu-N | 47,44 | N | | | |
| 2-Leu-CO | 169,954 | C | | | |
| 2-Leu-α | 52,197 | CH | 4,425 | 1,68 (β1), 1,53 (β2) | 136,17 (2-Leu-N), 176,86 (1-Leu-CO), 169,95 (2-Leu-CO), 35,86 (2-Leu-β), 71,34 (7-Lys-e) |
| 2-Leu-β | 35,859 | $CH_2$ | 1,680<br>1,534 | 4,43 (α), 1,53 (β2), 1,38 (γ)<br>4,43 (α), 1,68 (β2), 1,38 (γ) | 136,17 (2-Leu-N), 169,95 (2-Leu-CO)<br>169,95 (2-Leu-CO) |
| 2-Leu-γ | 24,469 | CH | 1,382 | 1,68 (β), 1,53 (β2), 0,87 (δ1), 0.82 (δ2) | 35,86 (2-Leu-β) |
| 2-Leu-δ1 | 22,722 | $CH_3$ | 0,868 | 1,38 (γ) | 35,86 (2-Leu-β) |
| 2-Leu-δ2 | 21,732 | $CH_3$ | 0,822 | 1,38 (γ) | 35,86 (2-Leu-β) |
| 2-Leu-N | 136,17 | N | | | |
| 3-Leu-CO | 170,520 | C | | | |
| 3-Leu-α | 50,530 | CH | 4,394 | 7,22 (NH), 1,38 (β1,2) | 169,95 (2-Leu-CO), 170,52 (3-Leu-CO), 41,82 (3-Leu-β) |
| 3-Leu-β | 41,823 | $CH_2$ | 1,384 | 4,39 (α), 1,49 (γ) | 118,74 (3-Leu-NH), 170,52 (3-Leu-CO) |
| 3-Leu-γ | 24,258 | CH | 1,490 | 1,38 (β1,2), 0,85 (δ1,2) | 41,82 (3-Leu-β) |
| 3-Leu-δ1 | 23,080 | $CH_3$ | 0,845 | 1,49 (γ) | 41,82 (3-Leu-β) |
| 3-Leu-δ2 | 21,698 | $CH_3$ | 0,848 | 1,49 (γ) | 41,82 (3-Leu-β) |
| 3-Leu-NH | 118,93 | NH | 7,22 | 4,39 (α) | 169,95 (2-Leu-CO) |
| 4-Ala-CO | 172,132 | C | | | |
| 4-Ala-α | 47,182 | CH | 4,429 | 8,22 (NH), 1,15 (β) | 170,52 (3-Leu-CO) |
| 4-Ala-β | 19,560 | $CH_3$ | 1,152 | 4,43 (α) | 121,54 (4-Ala-NH), 172,13 (4-Ala-CO) |
| 4-Ala-NH | 121,77 | NH | 8,22 | 4,43 (α) | 170,52 (3-Leu-CO) |
| 5-Leu-CO | 171,991 | C | | | |
| 5-Leu-α | 51,696 | CH | 4,099 | 8,07 (NH), 1,18 (β1), 1,12 (β2) | 40,56 (5-Leu-β) |
| 5-Leu-β | 40,555 | $CH_2$ | 1,179 | 4,10 (α), | 171.99 (5-Leu-CO) |

| | | | | | |
|---|---|---|---|---|---|
| | | | 1,113 | 4,10 (α) | 119,48 (5-Leu-NH) |
| 5-Leu-γ | 23,853 | CH | 1,171 | 0,76 (δ1), 0,71 (δ2) | 40,56 (5-Leu-β) |
| 5-Leu-δ1 | 22,507 | $CH_3$ | 0,760 | 1,17 (γ) | 40,56 (5-Leu-β) |
| 5-Leu-δ2 | 21,988 | $CH_3$ | 0,710 | 1,17 (γ) | 40,56 (5-Leu-β) |
| 5-Leu-NH | 119,49 | NH | 8,063 | 4,10 (α) | 172,13 (4-Ala-CO) |
| 6-Tyr-CO | 170,376 | C | | | |
| 6-Tyr-α | 55,897 | CH | 4,150 | 8,65 NH, 3,02 (β1), 2,61 (β2) | 171.99 (5-Leu-CO)<br>170,38 (6-Tyr-CO), 128,50 (6-Tyr-γ) |
| 6-Tyr-β | 35,779 | $CH_2$ | 3,023<br>2,610 | 4,15 (α), 2,61 (β2)<br>4,15(α), 3,02 (β1) | 119,59 (6-Tyr-NH), 170,38 (6-Tyr-CO), 128,50 (6-Tyr-γ)<br>119,59 (6-Tyr-NH), 170,38 (6-Tyr-CO), 128,50 (6-Tyr-γ) |
| 6-Tyr-γ | 128,497 | C | | | |
| 6-Tyr-δ,δ´ | 129,798 | CH | 7,025 | 6,63 (ε,ε´) | 155,74 (6-Tyr-ζ) |
| 6-Tyr-ε,ε´ | 114,793 | CH | 6,625 | 7,03 (δ,δ´) | 128,50 (6-Tyr-γ), 155,74 (6-Tyr-ζ) |
| 6-Tyr-ζ | 155,735 | C | | | |
| 6-Tyr-OH | | OH | 9,18 βr | | |
| 6-Tyr-NH | 119,80 | NH | 8,65 | 4,15 (α) | 171.99 (5-Leu-CO) |
| 7-Lys-CO | 175,078 | C | | | |
| 7-Lys-α | 53,009 | CH | 4,032 | 7,15 (NH), 1,67 (β1), 1,46 (β2) | 170,38 (6-Tyr-CO), 175,08 (7-Lys-CO) |
| 7-Lys-β | 31,658 | $CH_2$ | 1,667,<br>1,463 | 4,03 (α), 1.46 (β2), 1,07 (γ)<br>4,03 (α), 1,67 (β1), 1,07 (γ) | 71,34 (7-Lys-ε)<br>71,34 (7-Lys-ε) |
| 7-Lys-γ | 19,709 | $CH_2$ | 1,068 | 1,67 (β1), 1,46 (β2), 1,39 (δ1), 1,17 (δ2) | 71,34 (7-Lys-ε) |
| 7-Lys-δ | 33,749 | $CH_2$ | 1,385 | 1,17 (δ2), 1,07 (γ), 4,28 (ε)<br>1,39 (δ1), 1,07 (γ), 4,28 (ε) | 71,34 (7-Lys-ε) |

| | | | | | |
|---|---|---|---|---|---|
| | | | 1,168 | | |
| 7-Lys-ε | 71,342 | CH | 4,275 | 1,17 (δ2), 1,39 (δ1) | 47,44 (1-Leu-N) |
| 7-Lys-NH | 118,22 | NH | 7,15 | 4,03 (α) | 170,38 (6-Tyr-CO) |

Tab. S12d: Observed homonuclear and heteronuclear couplings and the chemical shifts ($^{1}H$ and $^{13}C$: δ(TMS)=0.00 ppm, $^{15}N$: δ($NH_3$)=0.00 ppm) of Apiosporin A3 in DMSO-$d_6$, at 308 K.
:

| position | δ ($^{13}C$,$^{15}N$) [ppm] | m ($^{13}C$, $^{15}N$) | δ ($^{1}H$) [ppm] | $J_{HH}$ | $^{n}J_{XH}$ |
|---|---|---|---|---|---|
| 1-Leu CO | 174,92 | C | | | |
| 1-Leu-α | 55,642 | CH | 3,275 | 1,49 (β1), 1,22 (β2) | 174,92 (1-Leu-CO) |
| 1-Leu-β | 40,475 | $CH_2$ | 1,491<br>1,219 | 3,28 (α), 1,81 (γ)<br>3,28 (α), 1,81 (γ) | 174,92 (1-Leu-CO), 24,53 (1-Leu-γ)<br>174,92 (1-Leu-CO), 24,53 (1-Leu-γ) |
| 1-Leu-γ | 24,521 | CH | 1,807 | 1,49 (β1), 1,22 (β2), 0,89 (δ1), 0,87 (δ2) | |
| 1-Leu-δ1 | 23,137 | $CH_3$ | 0,890 | 1,81 (γ) | 40,48 (1-Leu-β), 24,53 (1-Leu-γ)<br>21,62 (1-Leu-δ2) |
| 1-Leu-δ2 | 21,623 | $CH_3$ | 0,865 | 1,81 (γ) | 40,48 (1-Leu-β), 24,53 (1-Leu-γ)<br>23,14 (1-Leu-δ1) |
| 1-Leu-N | | N | | | |
| 2-Leu-CO | 168,99 | C | | | |
| 2-Leu-α | 54,102 | CH | 4,312 | 1,62 (β) | 168,99 (2-Leu-CO) |
| 2-Leu-β | 38,994 | $CH_2$ | 1,618 | 4,31 (α), 1,43 (γ) | |
| 2-Leu-γ | 24,706 | CH | 1,430 | 1,62 (β), 0,86 (δ1), 0,85 (δ2 | 22,28 (2-Leu-δ1) |
| 2-Leu-δ1 | 22,279 | $CH_3$ | 0,863 | 1,43 (γ) | 22,22 (2-Leu-δ2) |
| 2-Leu-δ2 | 22,223 | $CH_3$ | 0,852 | 1,43 (γ) | 22,28 (2-Leu-δ1), 24,70 (2-Leu-γ) , |
| 2-Leu-N | | N | | | |
| 3-Leu-CO | 171,02 | C | | | |

| | | | | | |
|---|---|---|---|---|---|
| 3-Leu-α | 50,580 | CH | 4,350 | 8,10 (NH), 1,48 (β1), 1,39 (β2) | 171,02 (3-Leu-CO) |
| 3-Leu-β | 40,045 | $CH_2$ | 1,465<br><br>1,397 | 4,35 (α), 1,40 (β2), 1,54 (γ)<br><br>4,35 (α), 1,47 (β1), 1,54 (γ) | 171,02 (3-Leu-CO), 22,94 (3-Leu-δ1), 21.21 (3-Leu-δ2)<br><br>171,02 (3-Leu-CO), 22,94 (3-Leu-δ1), ), 21.21 (3-Leu-δ2) |
| 3-Leu-γ | 24,154 | CH | 1,537 | 1,47 (β1), 0,86 (δ1), 0,82 (δ2) | |
| 3-Leu-δ1 | 22,944 | $CH_3$ | 0,855 | 1,54 (γ) | 24,15 (3-Leu-γ), 40,05 (3-Leu-β) |
| 3-Leu-δ2 | 21,206 | $CH_3$ | 0,815 | 1,54 (γ) | 24,15 (3-Leu-γ), 40,05 (3-Leu-β |
| 3-Leu-NH | 117,96 | NH | 8,100 | 4,35 (α) | 168,99 (2-Leu-CO) |
| 4-Ala-CO | 171,91 | C | | | |
| 4-Ala-α | 47,336 | CH | 4,383 | 8,02 (NH), 1,15 (β) | 171,91 (4-Ala-CO) |
| 4-Ala-β | 19,179 | $CH_3$ | 1,145 | 4,10 (α) | 171,91 (4-Ala-CO), 47,34 (4-Ala-α) |
| 4-Ala-NH | 120,76 | NH | 8,02 | 4,10 (α) | 171,91 (4-Ala-CO) |
| 5-Leu-CO | 171,58 | C | | | |
| 5-Leu-α | 51,315 | CH | 4,103 | 8,10 (NH), 1,16 (β1), 1,04 (β2) | 171,91 (4-Ala-CO) |
| 5-Leu-β | 40,752 | $CH_2$ | 1,171<br>1,048 | 4,10 (α), 1,05 (β2)<br>4,10 (α), 1,05 (β2) | 171,58 (5-Leu-CO)<br>171,58 (5-Leu-CO) |
| 5-Leu-γ | 23,777 | CH | 1,184 | 0,75 (δ19, 0,73 (δ2) | |
| 5-Leu-δ1 | 22,608 | $CH_3$ | 0,749 | 1,18 (γ) | |
| 5-Leu-δ2 | 21,759 | $CH_3$ | 0,730 | 1,18 (γ) | |
| 5-Leu-NH | 122,23 | NH | 8,10 | 4,10 (α) | 171,58 (5-Leu-CO) |
| 6-Tyr-CO | 171,16 | C | | | |
| 6-Tyr-α | 54,879 | CH | 4,301 | 8,26 (NH), 3,02 (β1), 2,61 (β2) | 171,16 (6-Tyr-CO) |
| 6-Tyr-β | 36,119 | $CH_2$ | 3,024<br>2,609 | 4,30 (α), 2,61 (β2)<br><br>4,30 (α), 3,02 (β1) | 129,78 (6-Tyr-δ)<br>127,79 (6-Tyr-γ), 129,78 (6-Tyr-δ) |
| 6-Tyr-γ | 127,793 | C | | | |

| | | | | | |
|---|---|---|---|---|---|
| 6-Tyr-δ,δ′ | 129,783 | CH | 6,996 | 6,62 (6-Tyr-ε) | 155,77 (6-Tyr-ζ) |
| 6-Tyr-ε,ε′ | 114,786 | CH | 6,624 | 7,00 (6-Tyr-δ) | 127,79 (6-Tyr-γ), 155,77 (6-Tyr-ζ) |
| 6-Tyr-ζ | 155,77 | C | | | |
| 6-Tyr-OH | | OH | n.δ. | | |
| 6-Tyr-NH | 118,00 | NH | 8,26 | 4,30 (α) | |
| 7-Lys-CO | n.δ | C | | | |
| 7-Lys-α | 52,494 | CH | 4,105 | 7,46 (NH), 1,70 (β1), 1,55 (β2) | |
| 7-Lys-β | 30,741 | $CH_2$ | 1,703<br>1,546 | 1,55 (β2), 4,11 (α)<br>4,11 (α), 1,70 (β1) | |
| 7-Lys-γ | 21,449 | $CH_2$ | 1,320<br>1,259 | | |
| 7-Lys-δ | 33,992 | $CH_2$ | 1,649<br>1,310 | 1,31 (δ2), 4,21 (ε)<br>1,65 (δ1), 4,21 (ε) | |
| 7-Lys-ε | 73,043 | CH | 4,213 | 1,65 (δ1), 1,31 (δ2) | |
| 7-Lys-NH | 114,91 | NH | 7,46 | 4,10 (α) | 171,16 (6-Tyr-CO) |

Tab. S12e: Observed homonuclear and heteronuclear couplings and the chemical shifts ($^{1}H$ and $^{13}C$: δ(TMS)=0.00 ppm, $^{15}N$: δ($NH_3$)=0.00 ppm) of Apiosporin A4 in DMSO-$d_6$, at 308 K.

| position | δ ($^{13}C$,$^{15}N$) [ppm] | m ($^{13}C$, $^{15}N$) | δ ($^{1}H$) [ppm] | $J_{HH}$ | $^{n}J_{XH}$ |
|---|---|---|---|---|---|
| 1-Leu CO | 176,47 | C | | | |
| 1-Leu-α | 56,62 | CH | 3,235 | 1,50 (β1), 1,36 (β2) | 176,47 (1-Leu-CO), 24,69 (1-Leu-γ) |
| 1-Leu-β | 41,17 | $CH_2$ | 1,503<br>1,358 | 1,36 (β2), 3,34 (α),1.79 (γ)<br>1,50 (β2), 3,34 (α),1.79 (γ) | 176,47 (1-Leu-CO), 24,69 (1-Leu-γ), 47,42 (1-Leu-N)<br>176,47 (1-Leu-CO), 24,69 (1-Leu-γ), 47,42 (1-Leu-N) |
| 1-Leu-γ | 24,69 | CH | 1,789 | 1,50 (β1), 1,34 (β2), 0,88 (δ1), 0,85 (δ2) | |
| 1-Leu-δ1 | 23,22 | $CH_3$ | 0,880 | 1,79 (γ) | 24,69 (1-Leu-γ) |

| | | | | | |
|---|---|---|---|---|---|
| 1-Leu-δ2 | 21,38 | $CH_3$ | 0,855 | 1,79 (γ) | 24,69 (1-Leu-γ) |
| 1-Leu-N | 47,42 | N | | | |
| 2-Leu-CO | 169,72 | C | | | |
| 2-Leu-α | 51,86 | CH | 4,471 | 1,68 (β1), 1,48 (β2) | 176,47 (1-Leu-CO), 69,72 (2-Leu-CO), 70,91 (7-Lys-e), 136,34 (2-Leu-N) |
| 2-Leu-β | 35,55 | $CH_2$ | 1,678<br>1,477 | 4,47 (α), 1,48 (β2), 1,38 (γ)<br>4,47 (α), 1,68 (β1), 1,38 (γ) | 169,72 (2-Leu-CO), 24,46 (2-Leu-γ), 136,34 (2-Leu-N)<br>169,72 (2-Leu-CO)<br>24,46 (2-Leu-γ) |
| 2-Leu-γ | 24,46 | CH | 1,383 | 1,68 (β1), 1,48 (β2), 0,87 (δ1), 0,82 (δ2) | |
| 2-Leu-δ1 | 22,52 | $CH_3$ | 0,871 | 1,38 (γ) | 24,46 (2-Leu-γ), 21,83 (2-Leu-δ2) |
| 2-Leu-δ2 | 21,83 | $CH_3$ | 0,823 | 1,38 (γ) | 24,46 (2-Leu-γ), 22,52 (2-Leu-δ1) |
| 2-Leu-N | 136,34 | N | | | |
| 3-Leu-CO | 170,56 | C | | | |
| 3-Leu-α | 50,36 | CH | 4,400 | 7,17 (NH), 1,38 (β1), 1,34 (β2) | 169,72 (2-Leu-CO)<br>172,56 (3-Leu-CO) |
| 3-Leu-β | 42,23 | $CH_2$ | 1,377<br>1,338 | 4,40 (α), 1,34 (β2), 1,51 (γ)<br>4,40 (α), 1,38 (β1), 1,51 (γ) | 24,24 (3-Leu-γ)<br>24,24 (3-Leu-γ) |
| 3-Leu-γ | 24,46 | CH | 1,509 | 1,38 (β1), 1,34 (β2), 0,87 (δ1), 0,85 (δ2) | |
| 3-Leu-δ1 | 21,65 | $CH_3$ | 0,865 | 1,51 (γ) | 24,24 (3-Leu-γ), 23,16 (3-Leu-δ2) |
| 3-Leu-δ2 | 23,16 | $CH_3$ | 0,847 | 1,51 (γ) | 24,24 (3-Leu-γ), 21,65 (3-Leu-δ1) |
| 3-Leu-NH | 118,965 | NH | 7,169 | 4,40 (α) | 169,72 (2-Leu-CO)<br>172,56 (3-Leu-CO) |
| 4-Ala-CO | 172,50 | C | | | |
| 4-Ala-α | 47,05 | CH | 4,431 | 8,26 (NH), 1,15 (β) | 172,56 (3-Leu-CO), 72,50 (4-Ala-CO), 121.61 (4-Ala-N) |

| | | | | | |
|---|---|---|---|---|---|
| 4-Ala-β | 19,50 | $CH_3$ | 1,150 | 4,43 (α) | 172,50 (4-Ala-CO), 121.61 (4-Ala-N) |
| 4-Ala-NH | 121,61 | NH | 8,264 | 4,43 (α) | 172,56 (3-Leu-CO) |
| 5-Leu-CO | 172,13 | C | | | |
| 5-Leu-α | 52,21 | CH | 3,967 | 8,11 (NH), 1,20 (β1), 1,15 (β2) | 172,50 (4-Ala-CO)<br>172.14 (5-Leu-CO) |
| 5-Leu-β | 40,20 | $CH_2$ | 1,200<br>1,144 | 1,14 (β2),1,10 (γ), 3,97 (α)<br>1,20 (β1),1,10 (γ), 3,97 (α) | 172.14 (5-Leu-CO), 23,64 (5-Leu-γ)<br>172.14 (5-Leu-CO) 23,64 (5-Leu-γ), 119,89 (5-Leu-N) |
| 5-Leu-γ | 23,64 | CH | 1,099 | 1,20 (β1), 1,14 (β2), 0,76 (δ1), 0,69 (δ2) | |
| 5-Leu-δ1 | 22,34 | $CH_3$ | 0,756 | 1,10 (γ) | 23,64 (5-Leu-γ) |
| 5-Leu-δ2 | 22,23 | $CH_3$ | 0,686 | 1,10 (γ) | 23,64 (5-Leu-γ) |
| 5-Leu-NH | 119,89 | NH | 8,115 | 3,97 (α) | 172,50 (4-Ala-CO),<br>172.14 (5-Leu-CO) |
| 6-Tyr-CO | 170,85 | C | | | |
| 6-Tyr-α | 55,42 | CH | 4,162 | 8,54 (NH), 3,10 (β1), 2,58 (β2) | 172.14 (5-Leu-CO)<br>170,85 (6-Tyr-CO)<br>128,31 (6-Tyr-γ) |
| 6-Tyr-β | 35,44 | $CH_2$ | 3,097<br>2,578 | 4,16 (α), 2,58 (β2)<br>4,16 (α), 3,10 (β1) | 170,85 (6-Tyr-CO), 128,31 (6-Tyr-γ), 119,68 (6-Tyr-N)<br>170,85 (6-Tyr-CO), 128,31 (6-Tyr-γ), 119,68 (6-Tyr-N) |
| 6-Tyr-γ | 128,31 | C | | | |
| 6-Tyr-δ,δ′ | 129,74 | CH | 7,022 | 6,63 (ε) | 155,72 (6-Tyr-ζ) |
| 6-Tyr-ε,ε′ | 114,78 | CH | 6,627 | 7,02 (δ) | 128,31 (6-Tyr-γ),<br>155,72 (6-Tyr-ζ) |
| 6-Tyr-ζ | 155,72 | C | | | |
| 6-Tyr-OH | | OH | | | 155,723 (6-Tyr-ζ) |

| | | | | | |
|---|---|---|---|---|---|
| 6-Tyr-NH | 119,68 | NH | 8,544 | 4,16 (α) | 172.14 (5-Leu-CO) |
| 7-Lys-CO | 173,45 | C | | | |
| 7-Lys-α | 51,24 | CH | 4,184 | 7,30 (NH), 1,64 (β) | 170,85 (6-Tyr-CO)<br>173,45 (7-Lys-CO) |
| 7-Lys-β | 30,39 | $CH_2$ | 1,641 | 4,18 (α), 1,05 (γ1), 0,96 (γ2) | 173,45 (7-Lys-CO) |
| 7-Lys-γ | 18,88 | $CH_2$ | 1,051<br>0,957 | 0,96 (γ2)1,64 (β),1,39 (δ1), 1,19 (δ2)<br>1,05 (γ1)1,64 (β),1,39 (δ1), 1,19 (δ2) | |
| 7-Lys-δ | 33,09 | $CH_2$ | 1,385<br>1,188 | 1,05 (γ1), 0,96 (γ2), 1,19 (δ2), 4,35 (ε)<br>1,05 (γ1), 0,96 (γ2), 1,39 (δ1), 4,35 (ε) | 70,91 (7-Lys-ε) |
| 7-Lys-ε | 70,91 | CH | 4,352 | 1,39 (δ1), 1,19 (δ2) | |
| 7-Lys-NH | 114,907 | NH | 7,303 | 4,18 (α) | 170,85 (6-Tyr-CO) |

Tab. S13: Ratio between L-/D- leucines enantiomers in Apiosporin A.

| | L | D | RATIOL/D |
|---|---|---|---|
| UV340 | 3136,6 | 1081,4 | 2,900499 |
| UV413 | 1224,5 | 418,8 | 2,92383 |

**Tab. S14**: Suggested molecular formulas of masses corresponding to lipids from monoisotopic masses of the *Ap. arundinis* AAU773 WT exudate droplet extracts.

| $[M+H]^+$ [Da] | Retention time [min] | Molecular formula | Δm [ppm] |
|---|---|---|---|
| 313.2738 | 15.10 | $C_{19}H_{36}O_3$ | -1.50 |
| 331.2839 | 15.10 | $C_{19}H_{38}O_4$ | -2.82 |
| 341.3059 | 16.05 | $C_{21}H_{40}O_3$ | 0.97 |
| 359.3160 | 16.05 | $C_{21}H_{42}O_4$ | -0.38 |

Tab. S15. The list of simulated cyclic peptides.

| Heptapeptides |
|---|

| | |
|---|---|
| 1 | cyclo(Leu-Leu(D)-Leu-Leu(D)-Leu-Tyr(D)-Lys) |
| 2 | cyclo(Leu-Leu(D)-Leu-Ala(D)-Leu-Tyr(D)-Lys) |
| Heptapeptides (L-conformers) | |
| 3 | cyclo(Leu-Leu-Leu-Leu-Leu-Tyr-Lys) |
| 4 | cyclo(Leu-Leu-Leu-Ala-Leu-Tyr-Lys) |
| Tetrapeptides | |
| 5 | cyclo(Tyr(D)-Tyr-Val(D)-Val) |
| 6 | cyclo(Trp(D)-Tyr-Val(D)-Val) |
| 7 | cyclo(Trp(D)-Phe-Val(D)-Val) |

Tab. S16. The list of simulated systems.

<table>
<tr><td></td><td>Number of peptides</td><td>Concentration of peptides in water</td><td>Initial simulation box size</td><td rowspan="7"></td></tr>
<tr><td rowspan="5">Simulation of peptide in water</td><td>1</td><td>3.3 mM</td><td rowspan="5">8.0 x 8.0 x 8.0 nm$^3$</td></tr>
<tr><td>2</td><td>6.7 mM</td></tr>
<tr><td>4</td><td>13.5 mM</td></tr>
<tr><td>8</td><td>26.9 mM</td></tr>
<tr><td>16</td><td>54.9 mM</td></tr>
<tr><td></td><td>Number of peptides</td><td>Initial concentration of peptides in water phase</td><td>Initial simulation box size</td><td>Number of dodecane molecules</td></tr>
</table>

| | | | | |
|---|---|---|---|---|
| Simulation of peptides | 4 | 13.5 mM | 8.0 x 8.0 x 16.0 nm$^3$ | 1030 |
